# Identification of rare-disease genes in diverse undiagnosed cases using whole blood transcriptome sequencing and large control cohorts

**DOI:** 10.1101/408492

**Authors:** Laure Frésard, Craig Smail, Kevin S. Smith, Nicole M. Ferraro, Nicole A. Teran, Kristin D. Kernohan, Devon Bonner, Xin Li, Shruti Marwaha, Zachary Zappala, Brunilda Balliu, Joe R. Davis, Boxiang Liu, Cameron J. Prybol, Jennefer N. Kohler, Diane B. Zastrow, Dianna G. Fisk, Megan E. Grove, Jean M. Davidson, Taila Hartley, Ruchi Joshi, Benjamin J. Strober, Sowmithri Utiramerur, Care4Rare Canada Consortium, Undiagnosed Diseases Network, Lars Lind, Erik Ingelsson, Alexis Battle, Gill Bejerano, Jonathan A. Bernstein, Euan A. Ashley, Kym M. Boycott, Jason D. Merker, Matthew T. Wheeler, Stephen B. Montgomery

## Abstract

RNA sequencing (RNA-seq) is a complementary approach for Mendelian disease diagnosis for patients in whom exome-sequencing is not informative. For both rare neuromuscular and mitochondrial disorders, its application has improved diagnostic rates. However, the generalizability of this approach to diverse Mendelian diseases has yet to be evaluated. We sequenced whole blood RNA from 56 cases with undiagnosed rare diseases spanning 11 diverse disease categories to evaluate the general application of RNA-seq to Mendelian disease diagnosis. We developed a robust approach to compare rare disease cases to existing large sets of RNA-seq controls (N=1,594 external and N=31 family-based controls) and demonstrated the substantial impacts of gene and variant filtering strategies on disease gene identification when combined with RNA-seq. Across our cohort, we observed that RNA-seq yields a 8.5% diagnostic rate. These diagnoses included diseases where blood would not intuitively reflect evidence of disease. We identified *RARS2* as an under-expression outlier containing compound heterozygous pathogenic variants for an individual exhibiting profound global developmental delay, seizures, microcephaly, hypotonia, and progressive scoliosis. We also identified a new splicing junction in *KCTD7* for an individual with global developmental delay, loss of milestones, tremors and seizures. Our study provides a broad evaluation of blood RNA-seq for the diagnosis of rare disease.

## Main

It is estimated that 350 million individuals worldwide suffer from rare diseases, which are for the most part caused by a mutation in a single gene [1–3]. Consequently, rare disease, when considered collectively, poses a major disease burden. The current overall molecular diagnostic rate in the clinical genetics setting is estimated at 50% [4]. Whole exome sequencing (WES) is among the most successful approaches for identifying causal genetic factors in rare disease [5–7]. The strategy consists in identifying rare (minor allelic frequency (MAF) ≤ 10^−3^), *de novo* or inherited variants with deleterious impact. For patients in whom exome-sequencing is not informative, RNA-seq has shown diagnostic utility in specific tissues and diseases [8–10]. This includes the application of RNA-seq on muscle biopsies in patients with genetically undiagnosed rare muscle disorders [8], and RNA-seq on cultured fibroblasts from patients with mitochondrial disorders [9]. RNA-seq of tissues is a very powerful strategy when such tissues are obtained as part of standard clinical care or are readily accessible. In many, if not most cases, biopsies are not performed for clinical care, and the tissues are difficult to access.

To assess RNA-seq from blood as general diagnostic tool, we sought to evaluate it for rare diseases of different pathophysiologies. We obtained RNA sequencing data from samples from 87 individuals, 56 affected by rare diseases and 31 unaffected family members (Fig. S1, Table S1). In total, 89.4% of patients were exome-negative. Patients represented 47 different diseases and were broadly classified into 11 distinct disease categories, with neurology, hematology and ophthalmology as the most frequent (Fig. 1A, Table S2). We integrated these data with RNA-seq data from healthy individuals from the Depression Genes and Network (DGN) cohort (N=909) [11], the Prospective Investigation of the Vasculature in Uppsala Seniors (PIVUS) project (N=65) [12] and the Genotype-Tissue Expression consortium (GTEx version 7) (N=620) cohorts [13] (Table S3).

**Figure 1:**
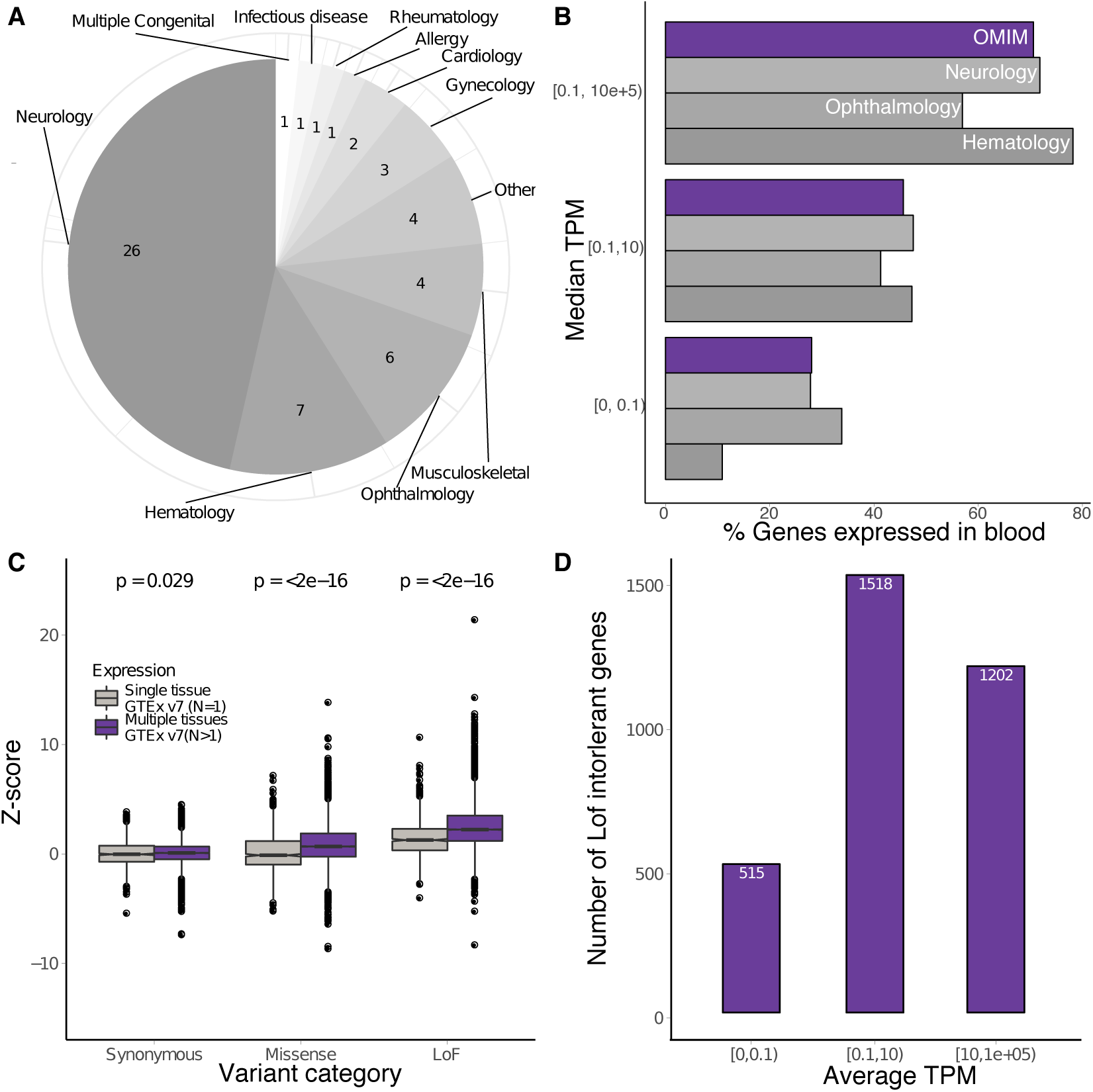
Using blood RNA-seq to study rare disease genes. A: Disease categories of sequenced affected patients. The majority of cases belong to neurology (n=26), hematology (n=7) and opthalmology (n=6) categories. B: Percentage of disease genes (from curated lists) expressed in blood. We used the median TPM across 909 DGN samples, 65 PIVUS samples and our 87 samples. We used disease categories that were the most represented in our dataset, in addition to OMIM genes. C: Tolerance to different types of mutations (from ExAC) in function of the expression status in a single versus multiple tissues. Genes that are expressed in multiple tissues tend to be more sensitive to missense and LoF mutations. D: Number of LoF intolerant genes stratified by expression level in blood. We considered genes with pLI score ≥ 0.9 as LoF intolerant.

## Results

### Blood expression in known disease genes

We first evaluated the extent that whole blood RNA-seq captured gene expression of known rare disease genes in each major disease category. We observed that the majority of known rare disease genes were expressed over 0.1 transcripts per million (TPM) (Fig. 1B, Table S4). When broadly considering disease genes from the Online Mendelian Inheritance in Man (OMIM) database [14], we observed 70.6% were expressed in blood and 50% of corresponding gene splicing junctions were covered with at least 5 reads in 20% of samples (Fig. S2). Notably, for a panel of genes known to be involved in neurological disorders (N=284), we observed that 72% were expressed. Using scores from ExAC [15], we further observed that genes expressed across multiple tissues were less tolerant to missense or loss-of-function (LoF) mutations (p-value ≤2 × 10^−16^), Fig. 1C, Table S3). This suggests that mutations that have more severe consequences occur more often in genes for which expression is not restricted to one tissue. Indeed, we observed that 66% of LoF-intolerant genes (probability of being intolerant to LoF mutations (pLI) ≥ 0.9) are expressed in blood samples (average TPM ≥ 1)(Fig. 1D).

### Gene expression outliers enriched for loss-of-function intolerance

Outlier (or aberrant) expression of a gene in a sample when comparing to all tested samples has previously been shown to help identify large-effect rare variants and rare disease genes in blood [16–18]. We assessed the differences between outlier genes in cases versus controls after correcting the data for batch effects (see Methods, Fig. S3, Fig. S4). We observed an enrichment of case outliers in genes more sensitive to LoF mutations (Fig. 2A, red, Fig. S5). This enrichment is more pronounced for under-expression outliers, which also corroborates the observations that new LoF mutations are more likely to lower expression level through nonsense mediated decay (NMD) [19–21]. As we increased the number of controls, the enrichment became stronger, demonstrating the importance of large control datasets to detecting outliers in LoF-intolerant genes (Fig. 2B, Fig. S6). We did not observe the same level of enrichment for missense sensitive genes and there was no enrichment of outliers for genes depleted in synonymous mutations.

**Figure 2:**
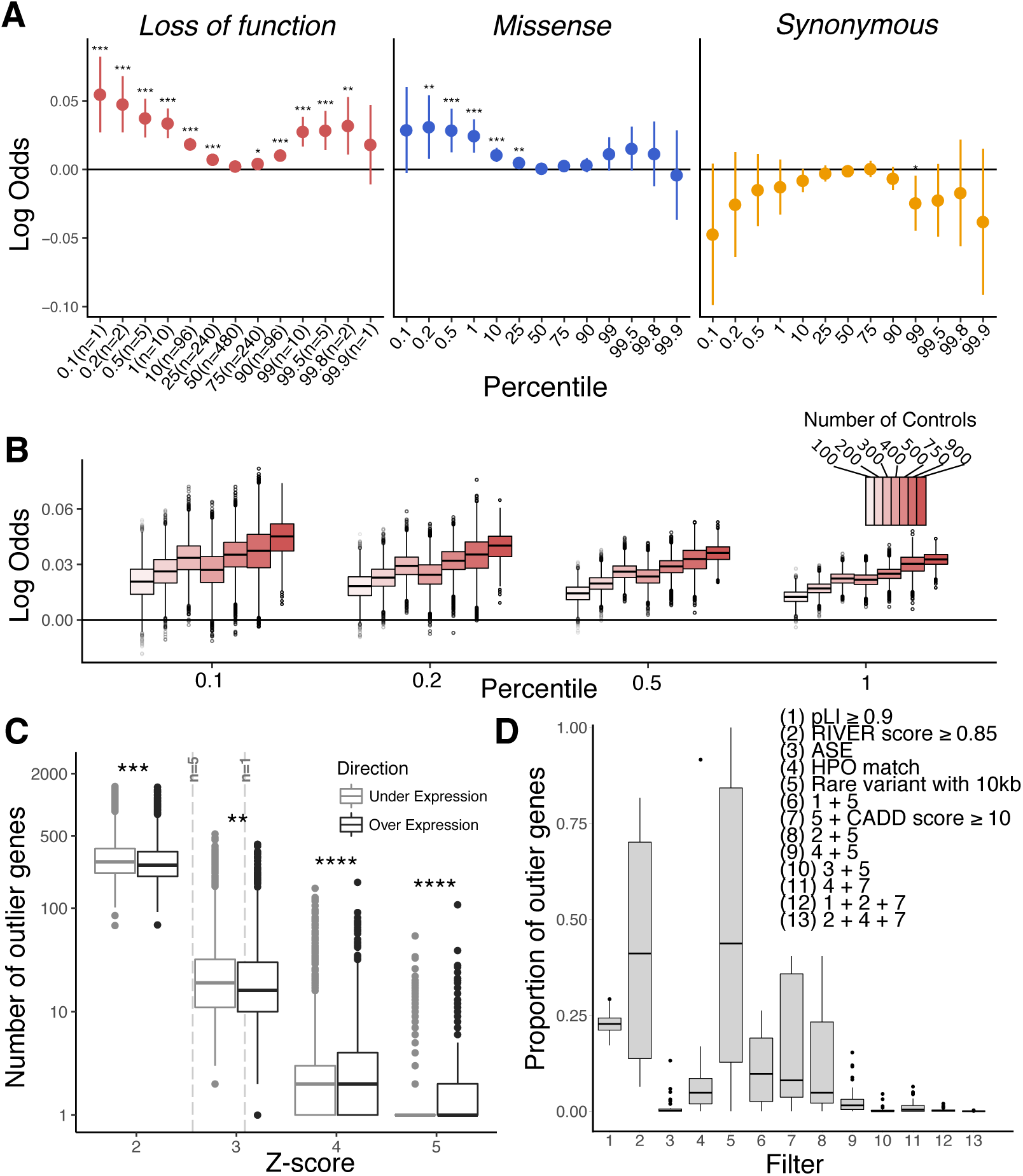
Expression outliers in rare disease samples. A: Enrichment for case or control outlier genes in intolerance to LoF (red), missense (blue) and synonymous (yellow) mutations at different percentiles of gene expression. B: Impact of the number of controls on enrichment in LoF model (10,000 permutations). Case enrichment increases as more controls are added. C: Impact of Z-score thresholds on number of outliers. Vertical dashed lines indicate mean Z-score for n=1 and n=5 percentiles across all genes used in analysis (N = 14,764). Significance level: **** p-value ≤ 1 ×10^−4^; *** p-value ≤ 1 ×10^−3^; ** p-value ≤ 1× 10^−2^; * p-value ≤ 5 ×10^−2^. D: Proportion of under-expression outlier genes remaining after filters. Adding genetic information (rare variant within 10kb of the gene body) allows to filter down to 50% of the original set of outliers. Keeping only genes for which HPO information of the affected individual match helps narrow down to less than 10% of candidates.

### Gene expression outliers in rare disease cases

We observed an average of 350 outliers per sample (|Z-score| ≥2, Fig. 2C, Fig. S7). At more extreme thresholds (|Z-score| ≥5), there are 1.2 outlier genes per sample. We tested different variant and gene-level filters that could aid in further narrowing down the lists of candidate genes (Fig. 2D). We filtered for genes that were LoF intolerant (Filter 1; pLI ≥ 0.9), likely to have a regulatory variant impacting gene expression (Filter 2: RIVER score ≥ 0.85); showed allele-specific expression (ASE) (Filter 3); linked to the phenotype (Filter 4: Human Phenotype Ontology [22] (HPO) match), with a rare variant with MAF ≤ 0.01% within 10kb upstream of the gene (Filter 5); and with a rare variant that was likely deleterious (Filter 7; CADD score ≥ 10). Other filters tested were combinations of these sets. We observed that when restricting to under-expression outlier genes with HPO matches and a deleterious rare variant nearby, we were able to reduce the candidate genes list to less than 10% of the initial set of outliers with 10% of cases having at least one candidate gene (Filter 11; Fig. S8A).

### Alternative splicing outliers in rare disease cases

Outlier splicing is also an important contributor to Mendelian disease [8, 9, 23–26]. To evaluate splicing events across rare disease samples, we corrected junction data for batch effects (Fig. S9) and obtained Z-scores in all samples (Fig. 3A, see Methods). On average, we detected 100 splicing outlier genes for each sample at |Z-score| ≥2 (Fig. 3B). We observed that the number of splicing outliers was influenced by the number of junction in each gene (Fig. S10), was higher in cases (Fig. S11) and, unlike expression outliers, was not enriched in genes sensitive to LoF or missense mutation (Fig. S12). From both exome and genome data alone, we observed that the number of candidate rare splicing variants was large but could be significantly reduced when combined with outlier splicing information from RNA-seq (Fig. 3C). From our pool of candidate genes with splicing outliers, we looked at the proportion remaining after different filters (Fig. 3D). We observed that limiting to genes relevant to the phenotype (Filter 2) and with a deleterious rare variant within 20 bp of the splicing junction (Filter 5), we were able to narrow down to only 0.05% of potential candidate genes (Filter 7). Overall, 15% of cases had at least one gene matching these criteria (Fig. S8B). Furthermore, genes selected after filtering carried more deleterious rare variants than unfiltered outliers suggesting an enrichment of disease-genes with compound heterozygous mutations (Fig. 3E).

**Figure 3:**
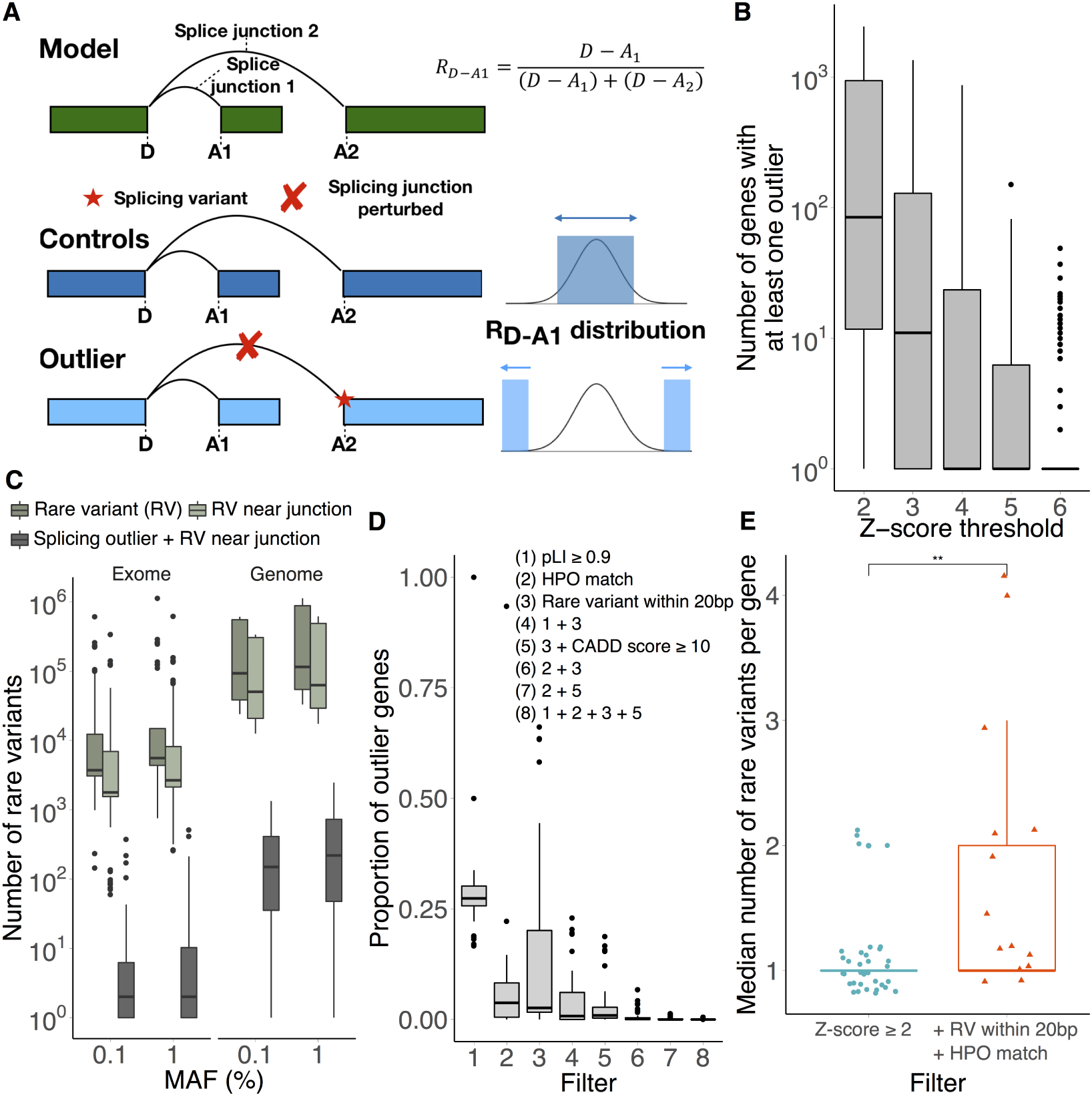
Splicing outlier detection. A: Splicing outlier definition. Gene model is in green, rectangles represent 3 exons. In this model, we show junction information for one donor (D) and two acceptors (A1 and A2). For each sample for this gene we have coverage information for the two existing splicing junctions (D-A1 and D-A2). We defined the proportion of one splice junction as the number of reads overlapping this junction divided by the total number of reads spanning all junction from a common donor (or acceptor). We calculated the proportion R of D-A1 through the equation on the right side of the figure. In the controls, no perturbation was observed in R. In case of a variant impacting splicing (red star), R in the affected sample will be very different from the other samples. B: Impact of the Z-score threshold on splicing outlier discovery. Number of genes with at least one splicing outlier at different Z-score thresholds. C: Number of rare variants in each sample, in total, nearby junction and associated with a splicing outlier. Rare variants were defined as variants with MAF ≤ 0.1%. D: Impact of different filters on splicing outlier discoveries. Combining gene and variant annotation with splicing outliers significantly reduced number of candidates. E: We observed a significant increase (Wilcoxon test, p-value 7.1× 10^−3^) in the median number of rare variants with CADD score ≥ 10 in the gene when filtering outliers with a rare variant within 20 bp of the junction and relevant to the disease phenotype (HPO match).

### Allele-specific expression outliers in rare disease cases

RNA-seq provides the ability to measure allele-specific expression. ASE can inform the presence of a large-effect heterozygous regulatory, splicing or nonsense variant, or epi-mutation aiding the identification of candidate rare disease genes and variants [9, 27–29]. Out of all possible heterozygous sites (*∼* 10^5^ to 10^6^ per sample for exome and genome, respectively), 10^4^ variants had sufficient coverage for analysis (Fig. 4A). Independent of sequencing technology, we observed 10^3^ sites displaying allelic imbalance with an allelic ratio ≤ 0.35 or ≥ 0.65. To highlight ASE events that might be disease-related, we focused on the subset of genes outlier ASE sites within case individuals when compared to all other rare disease individuals and GTEx samples (Fig. 4A). We found an average of 10 ASE outliers per individual. We next assessed the relevance of ASE outliers to case-specific phenotype-related genes as curated by Amelie. We observed that ASE outliers have significantly more genes with a non-zero Amelie score than random (Wilcoxon test p-value=0.015; Figure 4B). We also tested whether ASE would allow us to identify deleterious variants that were over-represented as this may be a marker for compound events or haploinsufficiency. Here, we focused on rare deleterious variants where the alternative allele is more abundant than the reference allele (Fig. 4C). In total, 25 rare variants show allelic imbalance biased toward the deleterious alternative allele (20 splice and 5 stop-gain). Among those, one variant is in *EFHD2*, a gene coding for Ca^2+^ adapter protein involved in B-cell apoptosis, NF-kB mediated inflammatory response, and immune cell activation and motility [30–33]. The carrier of this event was diagnosed with idiopathic cardiomyopathy, where accompanying symptoms (elevated inflammatory markers, Raynaud’s disease, and alopecia) are indicative of auto-immune issues.

**Figure 4:**
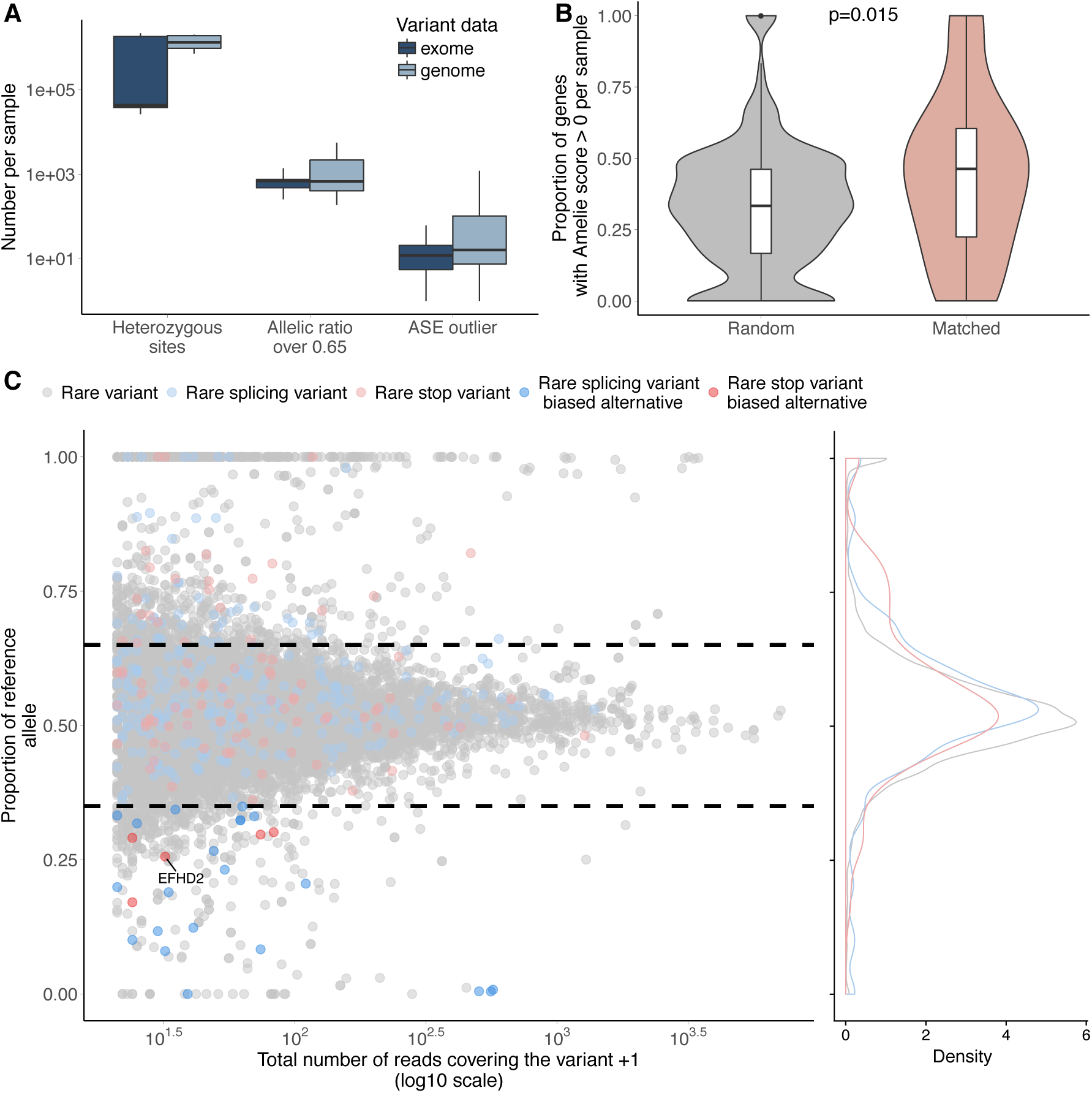
Allele specific expression across rare disease samples. A: Prevalence of ASE events. Results are displayed separately for exome and genome sequencing. B: Difference in proportion of non-zero Amelie scores for genes with outlier ASE in comparison to random genes (100 random gene sets for each sample). There are significantly more genes with non-zero Amelie scores in the non-random set (Wilcoxon rank test) indicating that the outlier ASE genes are influencing case-specific phenotype genes. C: Rare deleterious variants are biased towards the alternative allele across all samples. A stop-gain variant was highly expressed in *EFHD2* for one sample where there were matching symptoms.

### Diagnostic rate using blood RNA-sequencing

By integrating expression, splicing and ASE signals, we were able to identify and validate the causal gene in 4/47 independent cases (8.5%, 2 expression outliers, 2 splicing outliers), identify candidate genes potentially linked to the disease phenotype (gene matching HPO terms for the symptoms of the proband) in 9 cases (19.1%) and suggest novel genes not previously linked to the phenotype for 8 cases (17%, gene not matching HPO terms of the symptoms) (Fig. S14, Table S1). We did not find relevant candidate genes for 26 cases (55%). Notably, candidates were identified for three neurological cases where blood is not assumed to be a representative tissue.

### Expression outliers identify RARS2 casual gene

To represent how RNA-seq and subsequent filtering can enable disease gene discovery, we focused on one previously validated case. In this case, two sisters exhibited profound global developmental delay, neonatal-onset seizures, acquired microcephaly, hypotonia, G-/J-tube dependence, and progressive scoliosis and had undergone a diagnostic odyssey including comprehensive metabolic evaluation, storage disorder enzymology, and genetic testing. Exome sequencing identified two heterozygous pathogenic variants in the *RARS2* gene present in trans in both sisters: c.419T>G (p.F140C) and c.1612delA (p.T538fs). Pathogenic variants in *RARS2* had previously been associated with pontocerebellar hypoplasia type 6 (PCH6), a progressive neurodegenerative disorder reported in approximately 30 individuals in the literature to date [34–36]. The c.1612delA variant had never been reported in the literature in a patient with PCH6. RNA-seq identified *RARS2* as an under-expression outlier in one sister (Fig. 5) and with a Z-score=-1.55 in the other sister (Fig. S15). In the outlier sister, we were able to filter down from 1,724 candidate outlier genes to only 5 genes when limiting to those where a deleterious rare variant was within 10 kb of the gene body and had a HPO match. This is compared to 54 genes without any expression data. Among these five genes, *RARS2* was ranked first in terms of expression Z-scores.

**Figure 5:**
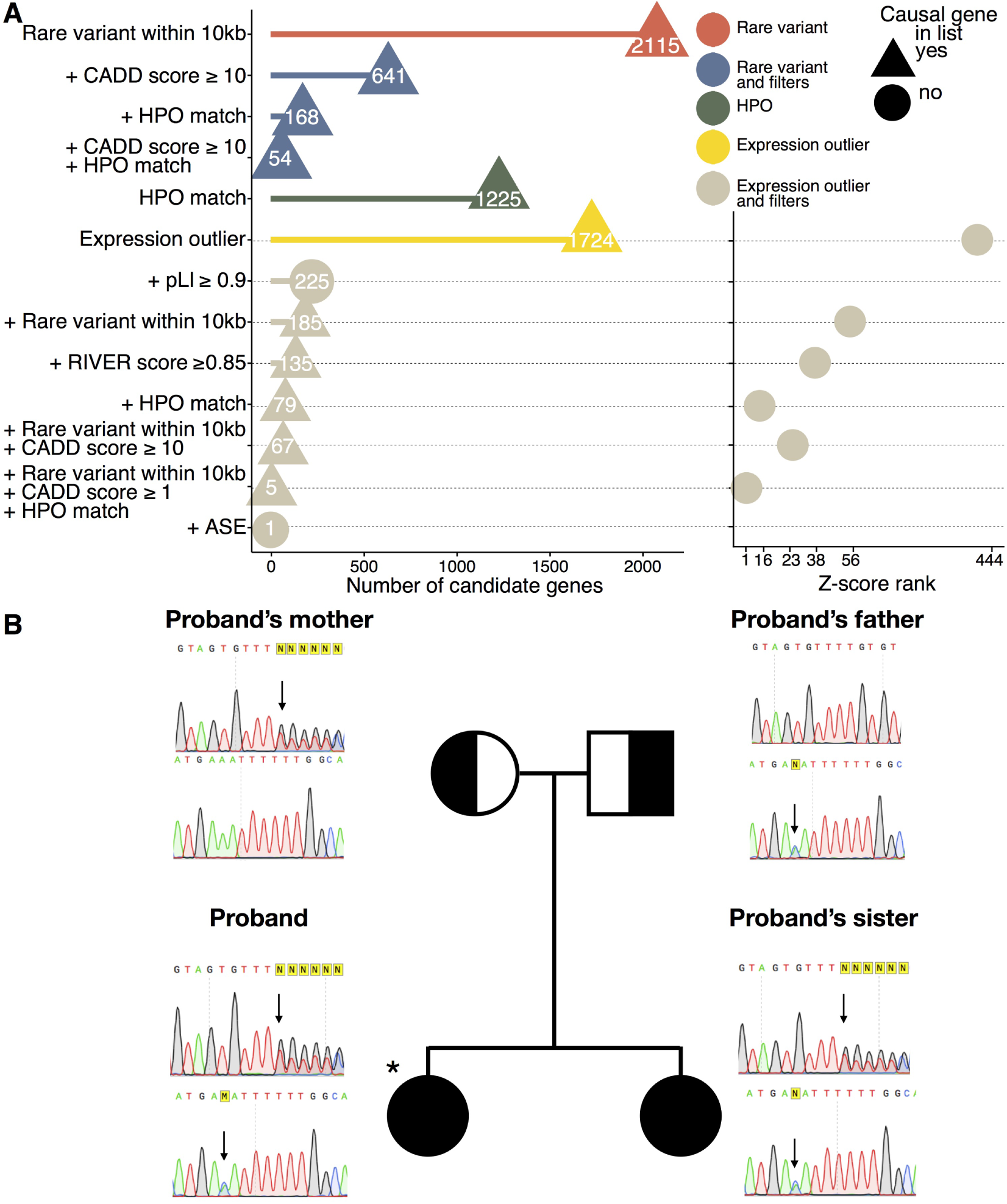
Identification of disease gene through expression outlier detection. A: Expression outliers in the *RARS2* case. Number of candidate genes obtained throughout different filters using genetic data, expression outlier data, and phenotypic data. Shape indicates if the causal gene is in the list. After filtering for expression outliers with a deleterious rare variant within 10 kb of gene body and limiting the search to genes for which there is a link to phenotype (HPO match), 5 candidate genes were left, with *RARS2* ranked first. B: Sanger sequencing showed c.1612delA and c.419T>G in the two affected sisters (proband marked with *) and their parents. Parents were carriers of one mutation and the two sisters carried both.

### Splicing outliers identify KCTD7 casual gene

To represent how spicing outliers and subsequent filtering can influence disease gene discovery, we focused on one previously unsolved case. In this case, a 12 year old Hispanic female presented with developmental regression after typical development until age 18 months, manifesting with loss of milestones including head control, and speech. Tremors developed at 21 months; and seizures at 22 months. She also suffered from occasional myoclonus. She has a 5-year-old brother with onset at 13 months of ataxia, autism, developmental delay, recurrent febrile seizures, and absent speech. We were able to filter the number of candidate genes from 2,543 genes to 33 genes, when looking only at genes associated with the phenotypes (from HPO terms [22]) and containing rare variants withing 20bp of annotated junctions with a CADD score ≥ 10 [37]. Adding splicing outlier information from RNA-seq data left us with two genes; *KCTD7*, containing a non-annotated junction in the affected sample, was the top-ranked (Fig. 6A, left panel). A synonymous mutation was found responsible for the creation of a new splicing junction in this gene (p.V152V, Fig. 6B). RT-PCR from RNA extracted from fibroblasts from exons 2-4 regions of the gene confirmed a difference in fragment size in the probands (Fig. 6C). In addition, this variant exhibited monoallelic expression towards the reference allele as a consequence of the premature splicing event (Fig. S16).

**Figure 6:**
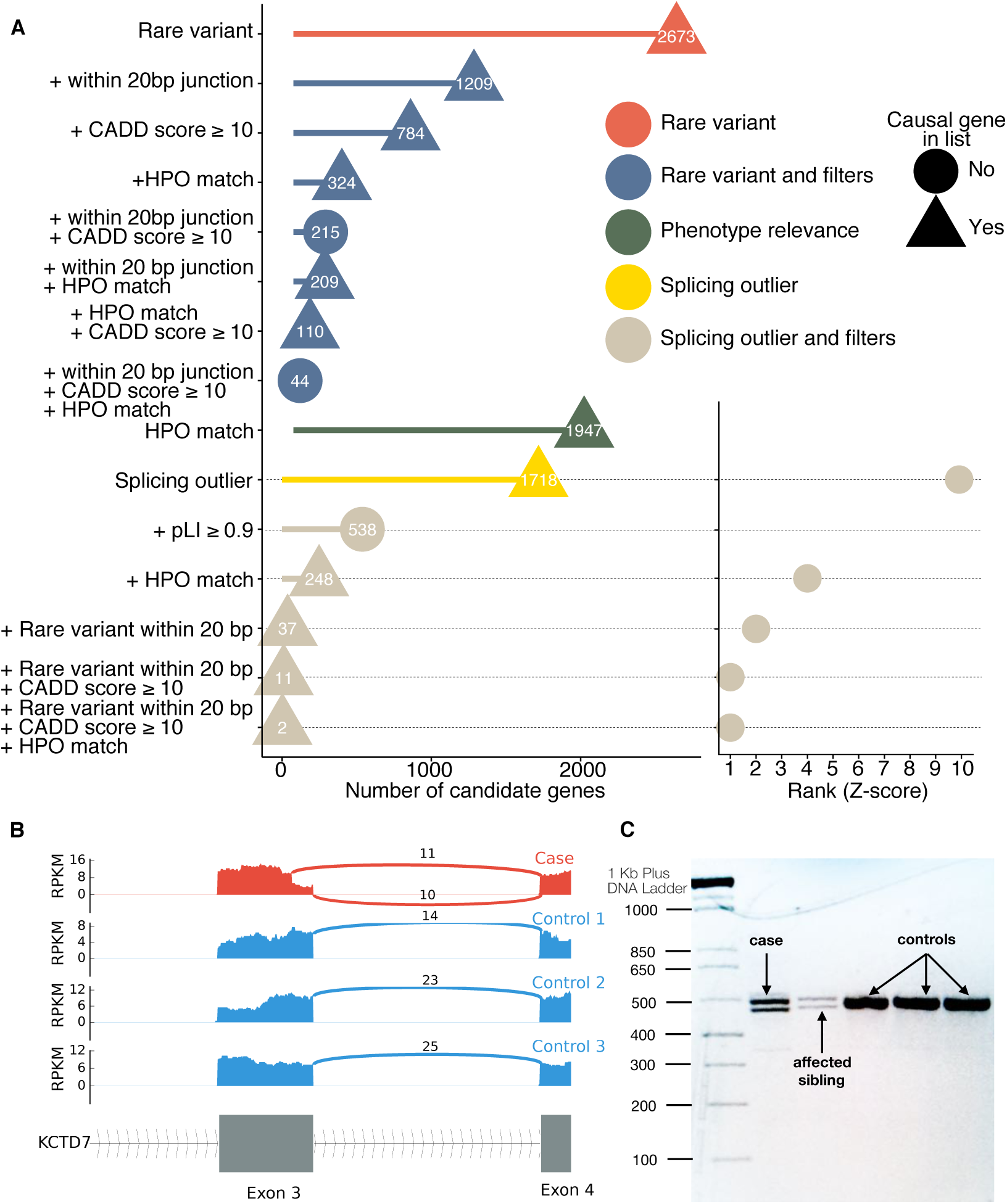
Identification of disease gene through splicing outlier detection. A: Splicing outliers in the *KCTD7* case. Number of candidate genes obtained throughout different filters using genetic data, splicing outlier data, and phenotypic data. Shape indicates if the causal gene is in the list. After filtering for splicing outliers with a deleterious rare splice variant within 20 bp of a splice junction and limiting the search to genes for which there is a link to phenotype (HPO match), 2 candidate genes were left, with *KCTD7* ranked first. B: Sashimi plot of the case and 3 controls of the splicing gain region in KCTD7. For the case only (red track), we observed a new splicing junction ahead of the annotated one in exon 3. C: cDNA gel from fibroblast cDNA of exons 2-4 of *KCTD7* for the proband, her affected sibling and three unaffected controls. Both for the case and her affected brother we observed 2 fragments of different size, corresponding to the alternative splice products induced by the splice-gain mutation. In control samples, only one fragment is observed, corresponding to the original transcript.

### Identification of causal disease genes using RNA-seq alone

While many of our cases had genetic data, use of RNA-seq alone can aid in disease gene identification. We reprocessed a solved case for which we had found an exon-skipping event in a previous study [10]. In this case, the patient presented with a sporadic form of spinal muscular atrophy. After filtering for splicing outliers (|Z-score| ≥ 2) and selecting only genes relevant to the symptoms (HPO), only one gene was left (Fig. S17A), corresponding to *ASAH* for which we subsequently identified with Sanger sequencing a splice-loss mutation leading to the creation of a new transcript, skipping exon 6 (Fig. S17B).

## Discussion

In summary, the use of whole blood RNA-seq in combination with variants and gene filters was able to identify the causal gene and variant(s) in 8.5% of cases or to highlight candidates genes linked to the phenotype in 19% of cases. This performance was demonstrated across a broad range of clinical cases with diverse clinical findings. Similar to the utility of large databases of control exomes for Mendelian disease diagnoses, we demonstrated the utility of large control RNA-seq data to identify aberrant expression, splicing and ASE events in candidate rare disease genes. Furthermore, this work demonstrates the utility of performing RNA-Seq on peripheral blood, which is a readily available specimen type in clinical practice. A trade-off needed to be found between strictly filtering the data and losing candidates of interest. We combined independent features of gene expression (aberrant expression level, splicing, ASE) to available data (variants, phenotypes) to prioritize genes of interest. It is worth noting that this combination of information is not expected to lead to the causal gene successfully in the following situations: first, if the causal gene is not expressed in the analyzed tissue; second, if the effects of the causal variant do not affect the expression of the gene; and third, if the filters are too strict (causal variant not present in exome-sequencing data, or gene not previously associated with the phenotype). Therefore, expert evaluation is still required when prioritizing candidate genes using RNA-seq data. We can expect that combining information from multiple ”omics” sources will only further improve diagnosis of unsolved rare disease cases in the future.

## Acknowledgments

SBM is supported by NIH grants R01HG008150 (NoVa) and U01HG009080 (GSPAC). LF was supported by the by the Stanford Center for Computational, Evolutionary, and Human Genomics Fellowship. CS is supported by BD2K Training Grant (T32 LM012409). NMF is supported by a National Science Foundation Graduate Research Fellowship. NAT is supported by the Stanford Genome Training Program (2T32HG000044-21). BL is supported by the Stanford Computational, Evolutionary and Human Genomics fellowship and the National Key R&D Program of China (2016YFD0400800). KMB is supported by a CIHR Foundation grant (FDN-154279). BB is supported by the Stanford Genome Training Program and Dean’s Postdoctoral Fellowship. CJP is supported by the NIST/JIMB grant 70NANB15H268. Clinical sample collection was supported, in part, by the Care4Rare Canada Consortium funded by Genome Canada, the Canadian Institutes of Health Research, the Ontario Genomics Institute, Ontario Research Fund, and Children’s Hospital of Eastern Ontario Foundation. Research reported in this manuscript was in part supported by the NIH Common Fund, through the Office of Strategic Coordination/Office of the NIH Director under Award Number U01HG007708. The content is solely the responsibility of the authors and does not necessarily represent the official views of the National Institutes of Health.

Members of the Undiagnosed Diseases Network; David R. Adams; Aaron Aday; Mercedes E. Alejandro; Patrick Allard; Euan A. Ashley; Mahshid S. Azamian; Carlos A. Bacino; Eva Baker; Ashok Balasubramanyam; Hayk Barseghyan; Gabriel F. Batzli; Alan H. Beggs; Babak Behnam; Hugo J. Bellen; Jonathan A. Bernstein; Gerard T. Berry; Anna Bican; David P. Bick; Camille L. Birch; Devon Bonner; Braden E. Boone; Bret L. Bostwick; Lauren C. Briere; Elly Brokamp; Donna M. Brown; Matthew Brush; Elizabeth A. Burke; Lindsay C. Burrage; Manish J. Butte; Shan Chen; Gary D. Clark; Terra R. Coakley; Joy D. Cogan; Heather A. Colley; Cynthia M. Cooper; Heidi Cope; William J. Craigen; Precilla D’Souza; Mariska Davids; Jean M. Davidson; Jyoti G. Dayal; Esteban C. Dell’Angelica; Shweta U. Dhar; Katrina M. Dipple; Laurel A. Donnell-Fink; Naghmeh Dorrani; Daniel C. Dorset; Emilie D. Douine; David D. Draper; Annika M. Dries; Laura Duncan; David J. Eckstein; Lisa T. Emrick; Christine M. Eng; Gregory M. Enns; Ascia Eskin; Cecilia Esteves; Tyra Estwick; Liliana Fernandez; Carlos Ferreira; Elizabeth L. Fieg; Paul G. Fisher; Brent L. Fogel; Noah D. Friedman; William A. Gahl; Emily Glanton; Rena A. Godfrey; Alica M. Goldman; David B. Goldstein; Sarah E. Gould; Jean-Philippe F. Gourdine; Catherine A. Groden; Andrea L. Gropman; Melissa Haendel; Rizwan Hamid; Neil A. Hanchard; Frances High; Ingrid A. Holm; Jason Hom; Ellen M. Howerton; Yong Huang; Fariha Jamal; Yong-hui Jiang; Jean M. Johnston; Angela L. Jones; Lefkothea Karaviti; David M. Koeller; Isaac S. Kohane; Jennefer N. Kohler; Donna M. Krasnewich; Susan Korrick; Mary Koziura; Joel B. Krier; Jennifer E. Kyle; Seema R. Lalani; C. Christopher Lau; Jozef Lazar; Kimberly LeBlanc; Brendan H. Lee; Hane Lee; Shawn E. Levy; Richard A. Lewis; Sharyn A. Lincoln; Sandra K. Loo; Joseph Loscalzo; Richard L. Maas; Ellen F. Macnamara; Calum A. MacRae; Valerie V. Maduro; Marta M. Majcherska; May Christine V. Malicdan; Laura A. Mamounas; Teri A. Manolio; Thomas C. Markello; Ronit Marom; Martin G. Martin; Julian A. Martínez-Agosto; Shruti Marwaha; Thomas May; Allyn McConkie-Rosell; Colleen E. McCormack; Alexa T. McCray; Jason D. Merker; Thomas O. Metz; Matthew Might; Paolo M. Moretti; Marie Morimoto; John J. Mulvihill; David R. Murdock; Jennifer L. Murphy; Donna M. Muzny; Michele E. Nehrebecky; Stan F. Nelson; J. Scott Newberry; John H. Newman; Sarah K. Nicholas; Donna Novacic; Jordan S. Orange; James P. Orengo; J. Carl Pallais; Christina GS. Palmer; Jeanette C. Papp; Neil H. Parker; Loren DM. Pena; John A. Phillips III; Jennifer E. Posey; John H. Postlethwait; Lorraine Potocki; Barbara N. Pusey; Genecee Renteria; Chloe M. Reuter; Lynette Rives; Amy K. Robertson; Lance H. Rodan; Jill A. Rosenfeld; Jacinda B. Sampson; Susan L. Samson; Kelly Schoch; Daryl A. Scott; Lisa Shakachite; Prashant Sharma; Vandana Shashi; Rebecca Signer; Edwin K. Silverman; Janet S. Sinsheimer; Kevin S. Smith; Rebecca C. Spillmann; Joan M. Stoler; Nicholas Stong; Jennifer A. Sullivan; David A. Sweetser; Queenie K.-G. Tan; Cynthia J. Tifft; Camilo Toro; Alyssa A. Tran; Tiina K. Urv; Eric Vilain; Tiphanie P. Vogel; Daryl M. Waggott; Colleen E. Wahl; Nicole M. Walley; Chris A. Walsh; Melissa Walker; Jijun Wan; Michael F. Wangler; Patricia A. Ward; Katrina M. Waters; Bobbie-Jo M. Webb-Robertson; Monte Westerfield; Matthew T. Wheeler; Anastasia L. Wise; Lynne A. Wolfe; Elizabeth A. Worthey; Shinya Yamamoto; John Yang; Yaping Yang; Amanda J. Yoon; Guoyun Yu; Diane B. Zastrow; Chunli Zhao; Allison Zheng

## Author Contributions

SBM, MTW, JDM, EAA and KMB conceived and planned the experiments. KSM, DB, JNK, DZ, DGF, MEG, JMD contributed to sample preparation. LL and EI provided phenotypic data together with blood RNA-seq of PIVUS control samples. SM, XL, KK, RJ, SU helped processing the variant data. LF carried out the analyses with the help of CS, NMF, NAT, ZZ, XL, BB, JRD, BL. KDK, BJS, AB, GB and JAB contributed to the interpretation of the results. KDK, CJP, DB, JNK, DZ, DGF, MEG performed the validation of results. LF and SBM wrote the manuscript with support from CS, NMF and NAT. All authors provided critical feedback and helped shape the research, analysis and manuscript.

## Material and Methods

### Rare disease cases and controls

We sequenced 87 whole blood samples, 56 extracted from affected individuals and 31 unaffected family members. The 56 cases represent a total of 47 independent diseases. Those samples were collected from 3 different institutions, the Children’s Hospital of Eastern Ontario (CHEO), the Stanford Clinical Genomics Program (CGP) and the Undiagnosed Disease Network (UDN).

Whole blood samples were sent to our lab in Paxgene RNA tubes or as isolated RNA for processing. Paxgene RNA tubes were processed manually per manufacturer’s protocol and 1.0 *µ*g RNA was used for further processing. Isolated total RNA was analyzed on an Agilent Bioanalyzer 2100 by pico RNA chip for RIN quality. Globin mRNA was removed using GLOBINclear prior to cDNA library construction. cDNA libraries were constructed following the Illumina TrueSeq Stranded mRNA Sample Prep Kit protocol and dual indexed. The average size and quality of each cDNA library was determined by Bioanalyzer and concentrations were determined by Qubit for proper dilutions and balancing across samples. On average, twenty pooled samples were run simultaneously on an Illumina NextSeq 500 (high output cartridge). Pooled samples were run in 6 distinct sequencing runs: two runs generated 75bp paired end reads and three runs generated 150 bp paired end reads. Output bcl files were converted to fastq files and demultiplexed using bcl2fastq version 2.15.0.4 from Illumina.

Reads were trimmed and adapters were removed using cutadapt (https://github.com/marcelm/cutadapt). Reads were then aligned to the reference human genome (hg19) with STAR v2.4.0j (https://github.com/alexdobin/STAR/releases/tag/STAR_2.4.0j). We used gencode v19 for reference annotation (https://www.gencodegenes.org/releases/19.html). We removed reads with a mapping quality under 30 and filtered duplicate reads with Picard Tools (http://broadinstitute.github.io/picard/). Gene-level and transcript-level quantifications were generated with RSEM v1.2.21 [38](https://github.com/deweylab/RSEM/releases/tag/v1.2.21). Junctions files generated by STAR were filtered: to consider a junction, a minimum of 10 reads uniquely spanning was required.

### Independent control cohorts for expression, splicing and ASE analyses

We used whole blood transcriptome data of 909 samples from the DGN cohort [11] as well as 65 samples (age 70) from the PIVUS cohort [12] to serve as independent healthy controls for expression analysis and splicing respectively. DGN samples are single-end 50bp reads and PIVUS samples are 75bp paired-end reads. Sequences were aligned, quantified and filtered following the same protocol used for rare disease cases and controls. We determined outlier ASE events at the gene level per individual by comparing our data to 620 individuals in GTEx v7 [39] across 48 tissues. Allele-specific expression in GTEx was processed as in [40], and only sites with a minimum of 20 reads overlapping and not entirely mono-allelically expressed were analyzed.

### Disease gene lists

Disease gene lists for neurology (n=284 genes), ophthalmology (n=380 genes) and hematology (n=50 genes) disease categories were obtained from curators for genes of interest in regards of the disease (Table S4). We obtained OMIM genes list (n=3,766 genes) from https://omim.org/downloads/. Gene expression of disease genes in our samples was restricted to protein coding genes.

### Genetic data

Variant data was produced according to recommended protocols for exome or genome data. VCFs obtained from UDN were generated through the Hudson Alpha and Baylor pipelines. In short, DNA reads alignment was performed using BWA-mem v0.7.12 [41] and variant calling was made using GATK v3.3 [42]. For CGS samples, variant calling was performed using GATK v3.4. We filtered variants according to the following criteria from previous studies [15, 43]:

- Filter field is PASS
- At least 20 reads covering the position (DP field)
- Genotype quality greater than 20 (GQ field)
- Normalized Phred-scaled likelihoods of the predicted genotypes lower than 20 (PL field)
- 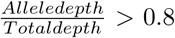 for homozygous calls and > 0.2 for each allele for heterozygous calls.
- Exclude variants with Hardy-Weinberg Equilibrium p-value < 1 × 10^−6^
- Exclude variants with call rate *<* 0.80 (missing > 20%)

We obtained genetic information for 75 samples (out of 87, 22 from whole genome sequencing, 53 from whole exome sequencing) (Fig. S13A). The number of LoF rare variants is variable across samples, and institutions (Fig. S13B). We merged all VCF files from those different institutions and homogenized their format for further analysis.

### Genetic data annotation

We annotated genetic data with allele frequency from the Genome Aggregation Database (gnomAD) [15] and Combined Annotation Dependent Depletion scores (CADD) [37] scores using Vcfanno (version 0.2.7) [44]. We used CADD scores v1.3 and gnomAD genomes release 2.0.2.

### Ancestry inference

VCF files were processed for ancestry inference using BCFtools v1.8 as following. They were normalized (fixing strand flips and left aligning indel records) and merged. We then subset this file to only variants in exonic regions, and filtered out variant with > 25% missingness. Missing variants were set to homozygous reference. A total of 11,016 variants remained after filtering. To perform ancestry inference, we used all individuals from 1000 Genomes phase 3 version 5 populations. For computational feasibility, we used genotypes from chromosomes 1,4,12,15,16, and 19. We used the prcomp function in R [45] to extract principal components and plotted the first three principal components.

### Expression levels normalization

We filtered out genes for which less than 50% of samples from each origin (i.e. rare disease individuals and unaffected family members sequenced in-house, external controls) had TPM > 0.5 and/or variance equal to zero. We performed Surrogate Variable Analysis (SVA) [46] using the ”two-step” method on a centered and scaled matrix of *log*_10_-transformed (*log*_10_(TPM+1)) RNA-seq count data output by RSEM [38]. We did not provide any known covariates to SVA. We added regression splines for Surrogate Variables (SVs) significantly associated with batch and/or institution (p-value < 1e-30 from linear regression of known covariates against all significant SVs). Linear regression splines had knots positioned at every 1.66% of samples, resulting in approximately 16 individuals per region - which is around the average number of individuals in each batch sequenced in-house (Fig. S4). Significant surrogate variables (SVs) and regression splines were then used as covariates in a regression model. The residuals of this model were centered and scaled to generate Z-scores for use in all subsequent analyses using gene expression data.

### Global outliers

To control for potential residual technical artifacts impacting outlier expression, we removed samples for which 100 or more genes had normalized expression values of |Z-score| > 4 after SVA correction (29 samples). We tested the model described in Figure 2 for several global outlier thresholds and observed a similar enrichment profile.

### Gene expression outlier enrichment analyses

In this analysis, we use the union of DGN samples and healthy family members that passed the global outliers criteria as the control set. We assessed enrichment for case outliers at increasingly stringent percentiles of gene expression in genes intolerant to mutations using a logistic regression model. As features in this model we used ExAC gene constraint metrics for LoF, missense and synonymous mutations [15]. For each gene in the dataset that had ExAC gene constraint metrics (n genes = 10535), we calculated a binomial outcome variable corresponding to the proportion of case expression outliers found in each gene: *Y*_*i*_ ∼ *B*(*n*_*i*_, *p*_*i*_), where *n*_*i*_ is the number of outlier samples in gene_*i*_ at a given percentile tested (the number of ’trials’), and *p*_*i*_ is the proportion of case outlier samples (which can be thought of as the probability of ’success’ (or all outliers being case outliers) for gene_*i*_). Then we model the relationship between the observed proportion of cases for each gene, and the corresponding gene constraint Z-score from ExAC. Specifically, we want to find *Pr*(*Y*_*i*_ = *All*_*C*_ *ases*|*X*_*i*_), where *X*_*i*_ is the gene constraint Z-score for gene_*i*_. We assess the effect of *X* using logistic regression: *logit*(*p*(*X*)) = β_0_ + β_1_*X*. A positive β_1_ value indicates that a step change in constraint metric *X* (toward genes less tolerant to mutations) is associated with an increase in the log odds of *Y*_*i*_ = 1 (i.e. all outliers being case outliers). A separate model was fit for each mutation class. We reported results as the log odds (+/- 1.96*SE) associated with each feature for each percentile. P-values were calculated based on the *z* -statistic.

### Rare variants around genes

We used BEDtools (version 2.26.0-112-gd8c0fe4) [47] to annotate gene and with rare variants within 10 kb. We filtered for rare variants with minor allele frequency from gnomAD ≤ 0.1%. We kept the singletons in the analysis.

### Junctions coverage ratios

Reference junctions were derived from Gencode v19 annotation file on known protein coding genes. A total of 142,246 in 14,296 genes. For each junction donor (then acceptor), all possible acceptors (then donors) were screened in the samples junctions files. The distribution of reads spanning those junction sets was attested by calculating the set ratios (Fig. 3A). We restricted the analysis to junctions for which several acceptors (donors) were associated to one donor (acceptor). In total, 13,109 groups of junctions were generated that way. In total, 25,941 junctions in 5,437 protein coding genes across all samples fulfilled those criteria.

### Splicing data normalization

In this analysis, we use the union of PIVUS samples and healthy family members as the control set. To remove as much noise as possible, and to allow missing values imputation, we removed junctions for which there was no more than 30 samples with data in the junction group (≤ 20% of samples). We analyzed coverage ratios for a total of 23,619 junctions.

Missing values in junction coverage ratios were imputed using missMDA R package [48]. PCA analysis was then performed using prcomp R package [45]. We regressed out principal components (PCs) accounting for 95% of the variation in our imputed dataset (129 PCs). We then put back original existing missing values in the dataset and derived Z-scores used in the outlier analysis. We looked at the correlation pattern between the 10 first PCs and known covariates from our dataset (Fig. S9). In brief, PC1 is mainly separating the source of the data (UDN, CGS, CHEO or PIVUS). PC2 is highlighting differences between the first batch and the other batches. Overall, we observe some level of correlation between all known covariates and the PCs that are regressed out from the data. We are assuming that the corrected data is exempt from most of the potential effects of those covariates.

### Rare variants around splicing junctions

We used BEDtools (version 2.26.0-112-gd8c0fe4) [47] to filter junction with a rare variant within 20 bp of a tested splicing junction. We filtered for rare variants with minor allele frequency ≤ 0.1%. We kept the singletons in the analysis.

### Allele specific expression

ASEReadCounter [49] version 3.8-0-ge9d806836 from Genome Anaysis Tool Kit (GATK) [50] was run on single nucleotide variants from VCFs provided by the UDN, CHEO and CGS and the RNA-seq data we processed. Only sites with a minimum read depth of 10, mapping quality of 10 and base quality of 2 were integrated in the analysis. For a gene to be considered with ASE, we required that at least 5 samples had heterozygous sites in the gene, that the heterozygous site was covered by at least 20 reads for the individual with ASE with an allelic ratio ≥ 0.65 or ≤ 0.35. We eliminated total mono-allelic expression from the analysis (ie allelic ratio = 0 or 1).

To detect ASE outliers we restricted our analysis to sites and genes common to our samples and GTEx dataset, including 8,758 genes and 61,447 sites, subject to the same site filters above. We scaled the reference ratios for all sites within a gene across samples to obtain |Z-scores| per site. To summarize GTEx data per individual, we considered the maximum ratio (| 0.5–reference_ratio |) across all tissues for which the individual had data at that site. We called an individual an ASE outlier for a gene if it had either the most extreme positive or negative Z-score, with |Z-score| > 2. Finally, we scored the ASE outlier genes based on their phenotypic relevance per sample using Amelie [51]. We generated a background distribution of Amelie scores per sample by randomly selecting the same number of genes for which a case was an ASE outlier from the set of all genes tested for ASE across both datasets, and scoring those genes against the case’s HPO terms using Amelie. This was repeated for 100 iterations per sample.

## RIVER

RIVER (RNA-informed variant effect on regulation) is a hierarchical Bayesian model to infer rare variants of their regulatory effects. Compared with other variant scoring methods, RIVER takes the advantage of utilizing both genomic information and transcriptome information [40].

We used GTEx v7 whole genome sequencing and cross-tissue RNA-seq data as training data for the model. The trained model (with learned parameters) is subsequently applied on UDN data to predict effects of rare variants. The model uses rare variants and the genomic annotations at those variants as predictors, uses RNA status (for this case is outlier status based on total gene expression levels) as the target/response variable.

Rare variants here are defined as those of minor allele frequency (MAF) *<* 0.01 in 1000 genome project phase III all populations combined. For variants in GTEx we additionally require MAF *<* 0.01 within GTEx cohort itself and for variants in the rare disease samples we additionally require MAF < 0.02 within the rare disease samples themselves. We considered all rare variants 10kb near genes (10kb before transcription start site until 10kb after transcription end site). Overall, there were a median of 2 rare variants per gene and individual pair for GTEx subjects and rare disease subjects. For this analysis, we considered protein-coding and lincRNA genes only.

We used the following genomic annotations: Ensembl VEP [52], CADD [37], DANN [53], conservation score (Gerp [54], PhyloP [55], PhastCons [56]), CpG content, GC content, chromHMM [57] and Encode chromatin-openness track. We selected those features based on their prior evidence of association with regulatory effects [40]. Features were aggregated over each gene and individual pair, using either max(), min() for quantitative features, or any() for categorical features.

Expression outliers (the response variable) were defined as those with |Z-score| > 2. Z-scores were calculated based on total gene expression level RPKM from RNA-seq. In addition, for GTEx training data, gene expression levels were corrected by PEER [58] to remove technical artifacts and major common-variant eQTL effects are also removed. Z-scores for GTEx are median over all available tissues [40].

### Phenotypic data

For each case we have RNA-seq data for, we also obtained HPO terms corresponding to the symptoms of the affected individual. We extended this list of HPO terms to terms that were hierarchically one level lower (child terms), one level higher (parent terms) or alternative terms for the same phenotype. To do so, we used the Human Phenotypic Ontology (HPO) [22] (http://human-phenotype-ontology.github.io/downloads.html). To link HPO terms to genes we used the genes to phenotype and phenotype annotation files provided by the Human Phenotypic Ontology. We used Amelie [51] to rank expression and splicing outlier candidates together with genes with outlier ASE events in regards of the relatedness of the gene to the cases HPO terms.

### Diagnostic rate

We labeled ”solved” cases for which we found candidates from RNA-seq data for which the causal mutation was found and validated. ”Linked to phenotype” candidates correspond to cases for which we found a good candidate gene linked to the phenotype of the affected individual that has not been validated yet. ”No direct link to phenotype” refers to cases for which we found a gene not directly linked to the disease phenotype with aberrant expression pattern and for which further information is needed. Cases for which no strong candidate genes were found after analyzing RNA-seq data are labeled ”no candidate”.

## Supplemental Material

Table S1: Overview of clinical cases integrated in the study

Table S2: Categories used to describe disease type of each proband

Table S3: Cohorts used as controls in our analysis

Table S4. Disease gene lists used in the analysis

---

**Fig. S1:**
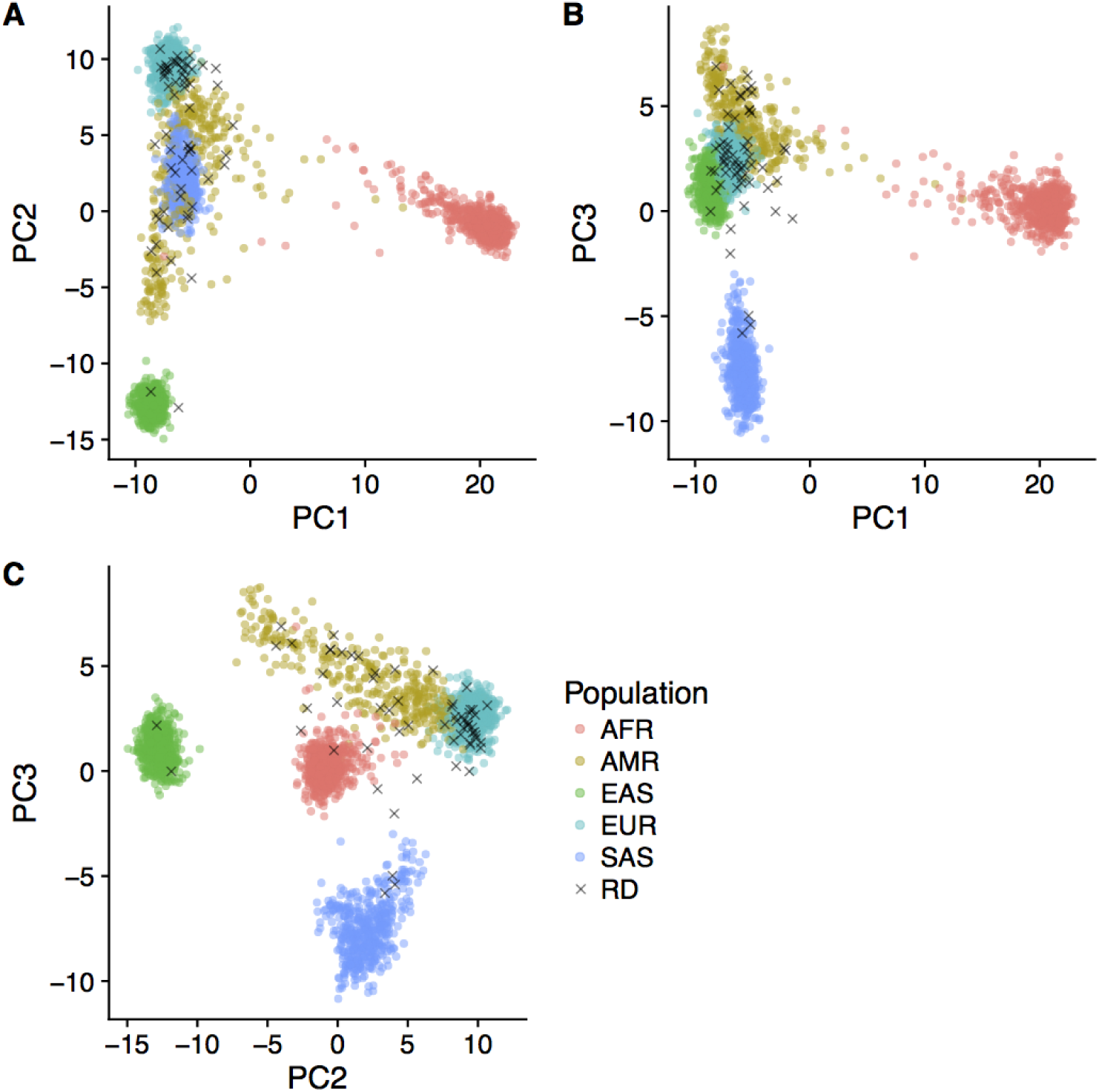
Ancestry principal components. We inferred ancestry principal components for all rare disease samples with genotype data using 1000 genome phase 3 version 5 as a reference. Panel (A) - (C) shows pairwise scatterplot for principal components 1 to 3. Abbreviation: RD = rare disease, AFR = African, AMR = Native American, EAS = East Asian, EUR = European, SAS = South Asian.

**Fig. S2:**
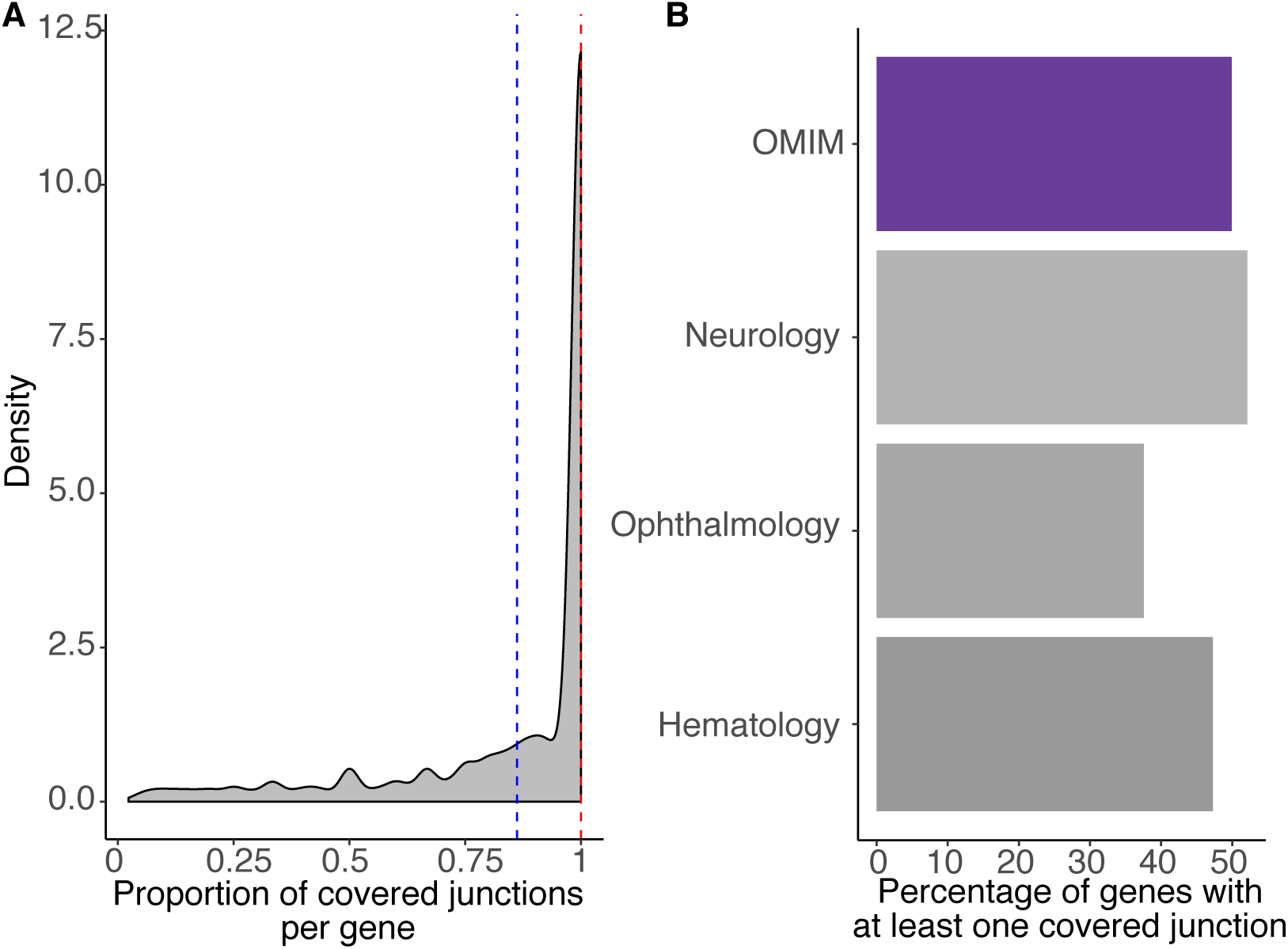
Junctions coverage across whole blood samples. We used a total of 1,061 whole blood samples from our controls cohorts and rare disease samples. A: Density plot representing the proportion of annotated junctions covered per gene. Those are a subset of genes for which at least one junction is covered with at least 5 uniquely mapped reads across at least 20% of the samples. On average (blue dashed line) 86%, (median of 100% - red dashed line) of junctions fulfill those criteria. B: Percentage of genes from disease genes panels in which at least one junction is covered with at least 5 uniquely mapped reads in at least 20% of samples. We observe that about 50% of genes from OMIM, Neurology, Ophthalmology or Hematology panels are fulfilling this criteria.

**Fig. S3:**
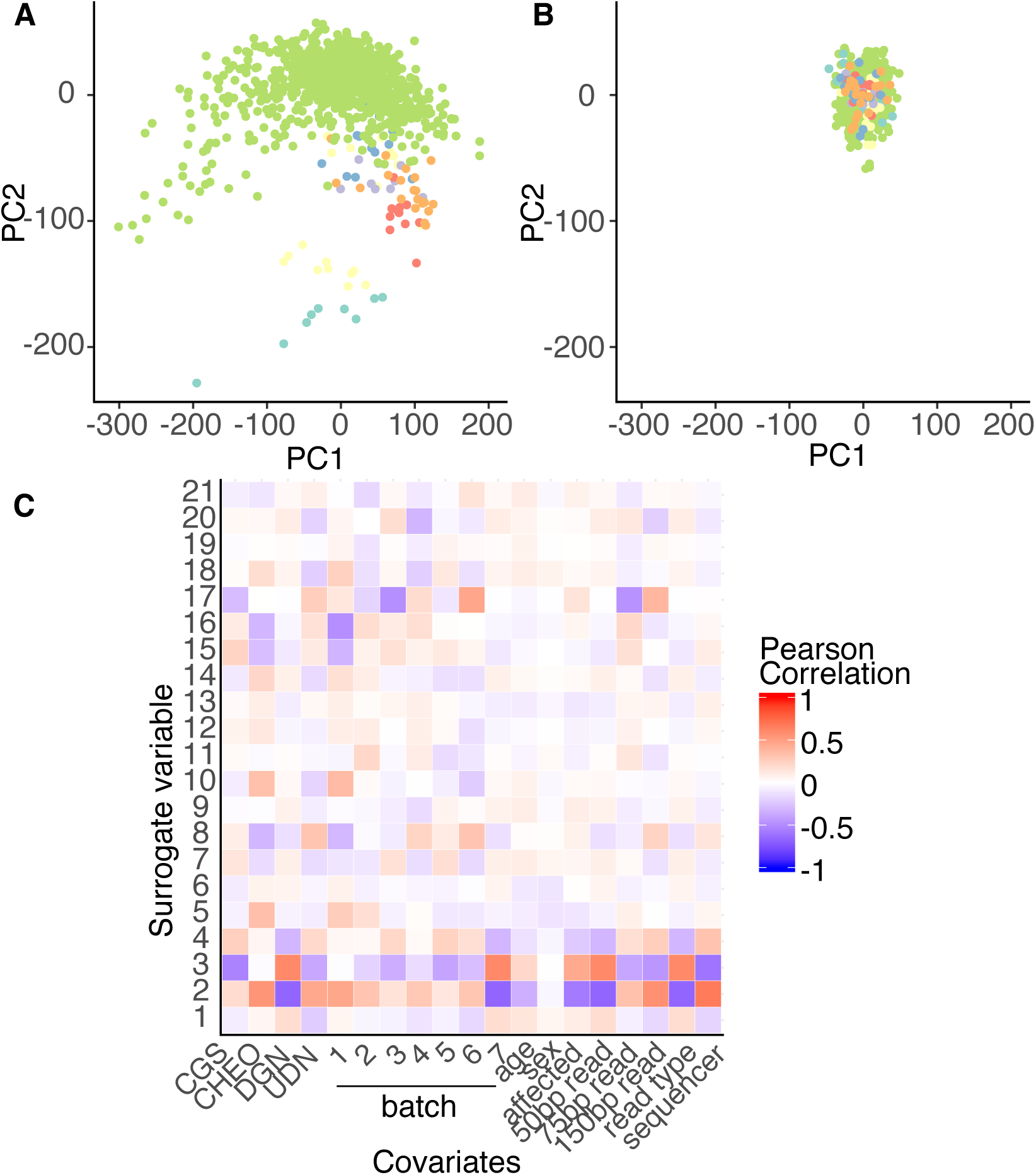
Correction for batch effects - Expression data. A: Plot of first two principal components run on uncorrected gene expression data. Samples are colored by batch. Largest cluster (green dots) is DGN control samples. B: Plot of first two principal components run on gene expression data after regressing out significant surrogate variables found by SVA. C: Correlation between known covariates and all significant surrogate variables (SVs). We did observe that SV2 is highly correlated with the read type, and the sequencing technology which corresponds to differences between DGN and the other samples. Overall, the first SVs are correcting for batch effects across our samples.

**Fig. S4:**
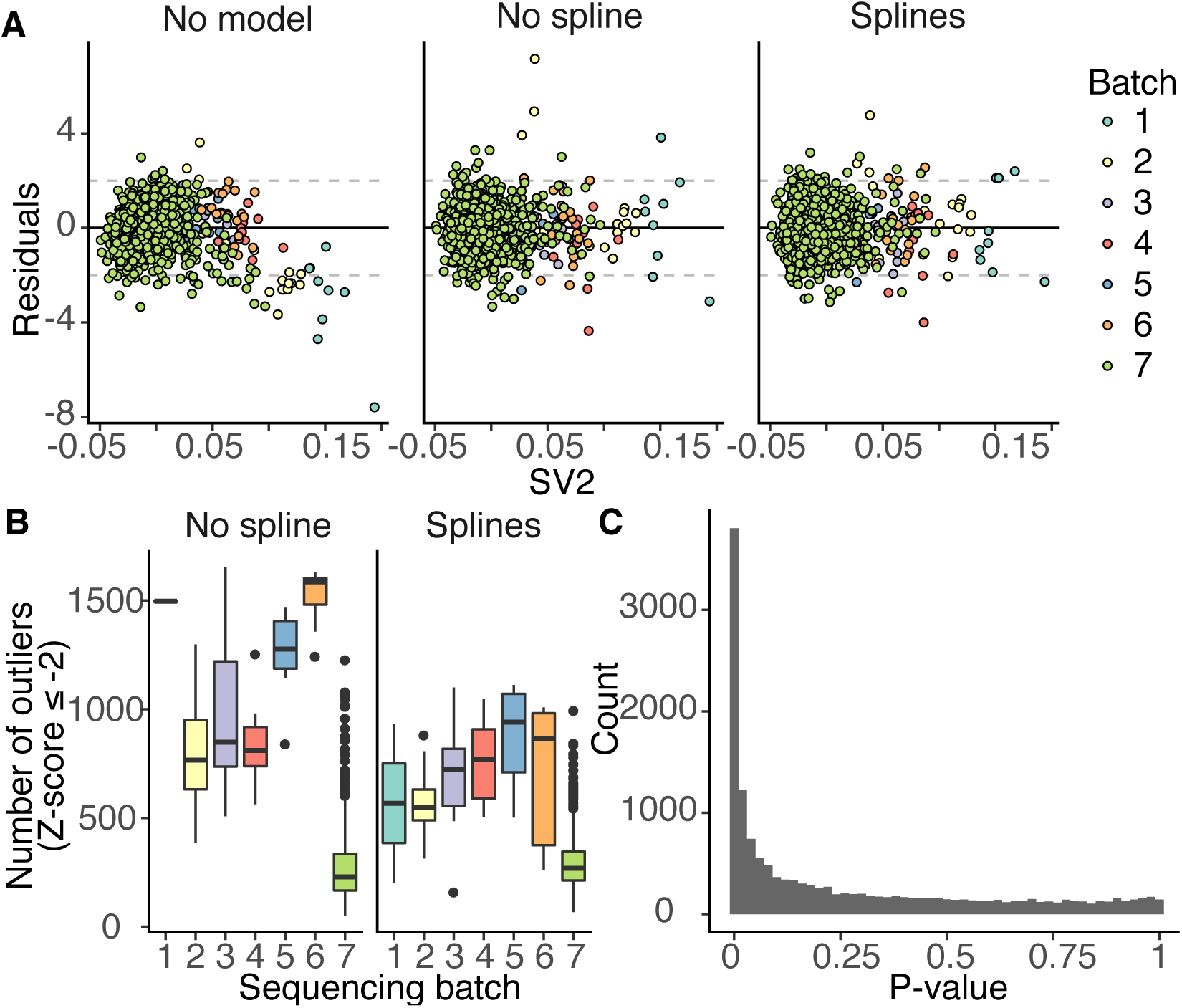
Use of regression splines in expression data. A: Normalized gene expression residuals, without correction (left panel), after regressing out significant surrogate variables (SVs) (middle panel) and significant SVs plus regression splines on SVs significantly associated with batch (right panel, p-value<1e-30). Residuals were plotted against SV2 - significantly associated with batch. Adding splines reduces the number of outliers without removing biologically-significant outliers. B: Number of outliers genes per batch (Z-score≤ -2) after correction with SVs (left panel) and SVs with regression splines on SVs associated with batch (right panel). Regression splines results in a more consistent number of outliers across all batches. C: Benjamini & Hochberg adjusted p-values resulting from a per-gene likelihood ratio test comparing linear regression model fit both with and without regression splines on SVs associated with batch and/or institution. Regression splines improve the model fit for 4,126 genes (p-value ≤ 0.05, 27.9% of all genes in dataset).

**Fig. S5:**
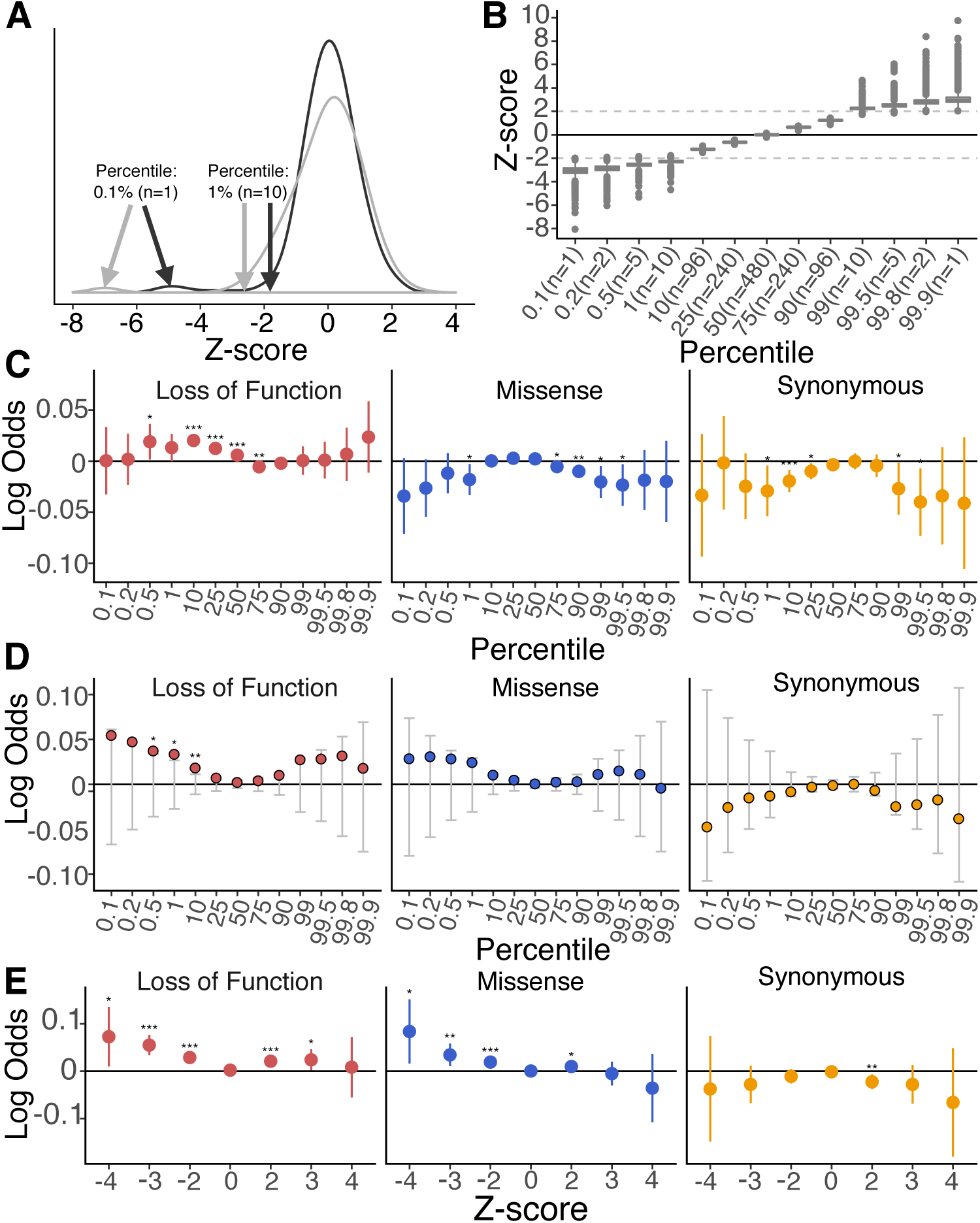
Further information on outlier enrichment analysis. A: Z-score may not correspond to expected gene expression distribution percentile due to outlier samples skewing the normal distribution. Finding fixed percentiles of samples ensures consistent sample sizes across all genes. B: Mapping between percentile and Z-score across thirteen under- and over-expression gene expression percentile thresholds. Percentiles are particularly useful for finding the least-expressed or most-expressed sample in a given gene. C: Based on Fig. 2A. Log odds testing proportion of case outliers found in mutation-intolerant genes. Case and control labels of rare disease individuals and unaffected family members have been switched, resulting in the previously observed enrichment no longer being evident. D: Observed log odds (color dots) relative to mean log odds (+/- 1.96*SE) resulting from 10,000 permutations of case/control labels across individuals. Asterisks indicate significance of observed log odds given permutation results (empirical p-value). E: Based on Fig. 2A but using Z-scores in place of percentiles. We observed a similar enrichment profile as found in Fig. 2A.

**Fig. S6:**
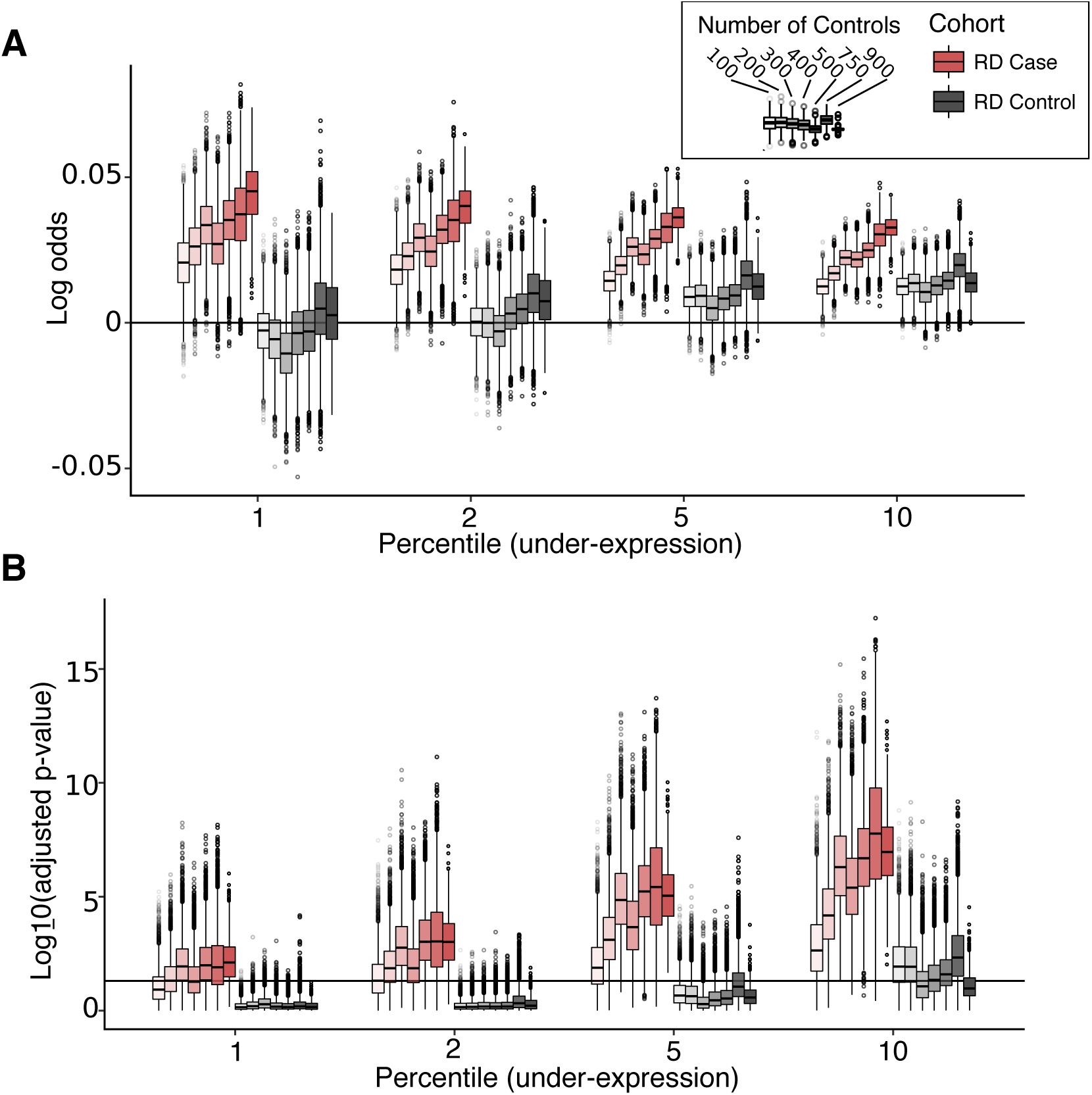
Impact of the number of controls on the enrichment. A: Enrichment of cases (red) under-expression outliers in LoF sensitive genes as we increase the number of controls (8,200 random subsets). This enrichment was not observed for rare disease family members controls (gray). B: Benjamini & Hochberg adjusted p-values associated with the enrichment at different number of controls. Horizontal line indicates 0.05 significance cutoff. The p-values are decreasing as we increase the number of controls. This observation is only true for case enrichment and holds at different under-expression percentile thresholds. When switching cases for controls (gray) we did not see the same pattern, particularly when using 909 external control samples.

**Fig. S7:**
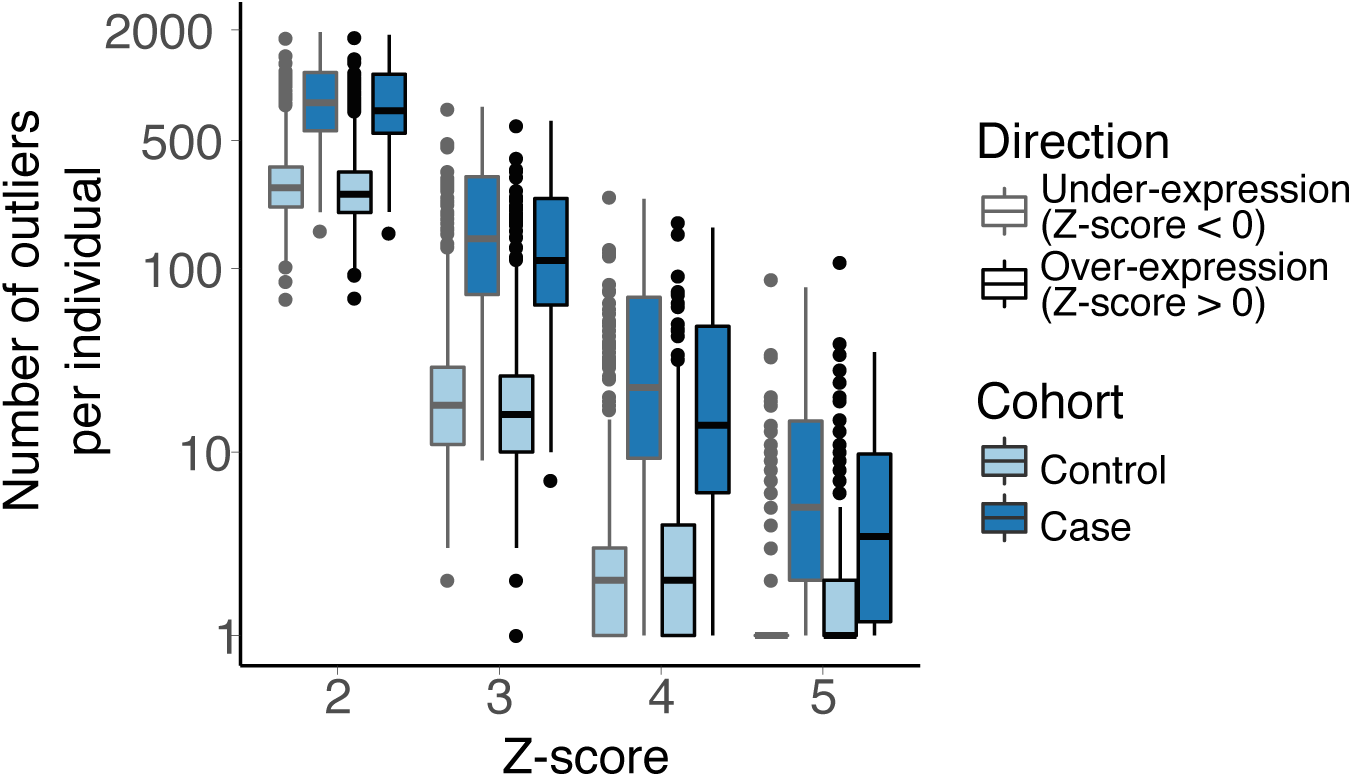
Impact of the affected status on number of outliers. Distribution of total number of outlier genes per sample across increasingly stringent Z-score thresholds for controls (light blue) and cases (dark blue) samples. Under-expression outliers are outlined in light gray, over-expression outliers are outlined in black. As Z-score threshold becomes more stringent, we observed fewer outlier genes per sample. There are more outlier genes in cases in comparison to controls at every Z-score cut-off for both under-expression and over-expression.

**Fig. S8:**
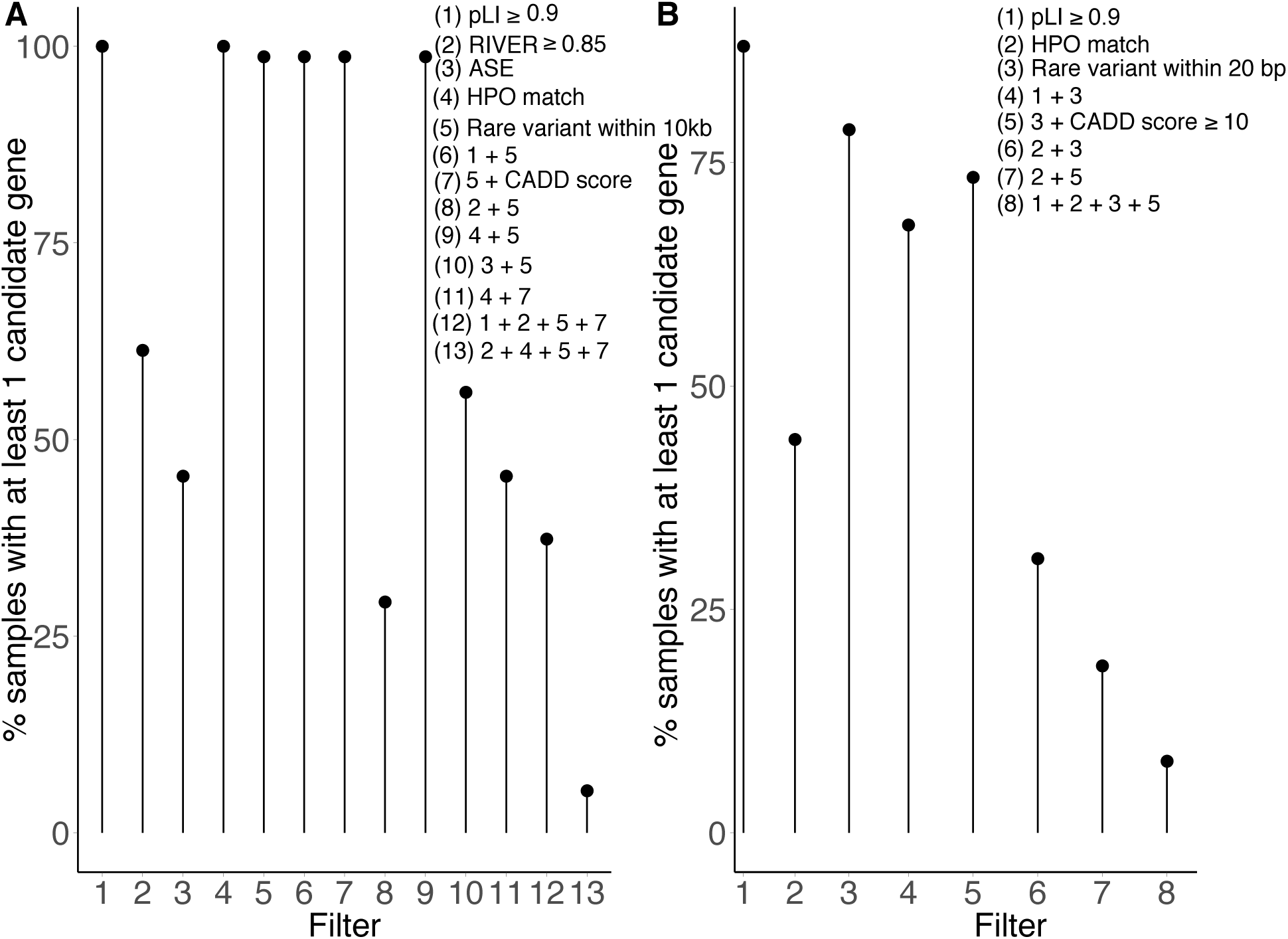
Percentage of samples with at least one candidate genes when filtering outliers. Filters have various impacts on the number of samples with at least one candidate gene. By combining several layers of filters we are drastically reducing the number of candidate genes but also the number of samples for which we have candidates. We recommend to relax filter stringency after looking at sets of genes that match the most stringent criterion. A: Expression outliers. After filtering for genes with a high RIVER score, matching HPO terms, with a deleterious rare variant within 10kb, we observed less than 10% of samples with at least one candidate gene (13). B: Splicing outliers. When keeping only genes with HPO match, and a deleterious rare variant with 20bp of the outlier junction, we observed candidate genes in less than 10% of samples.

**Fig. S9:**
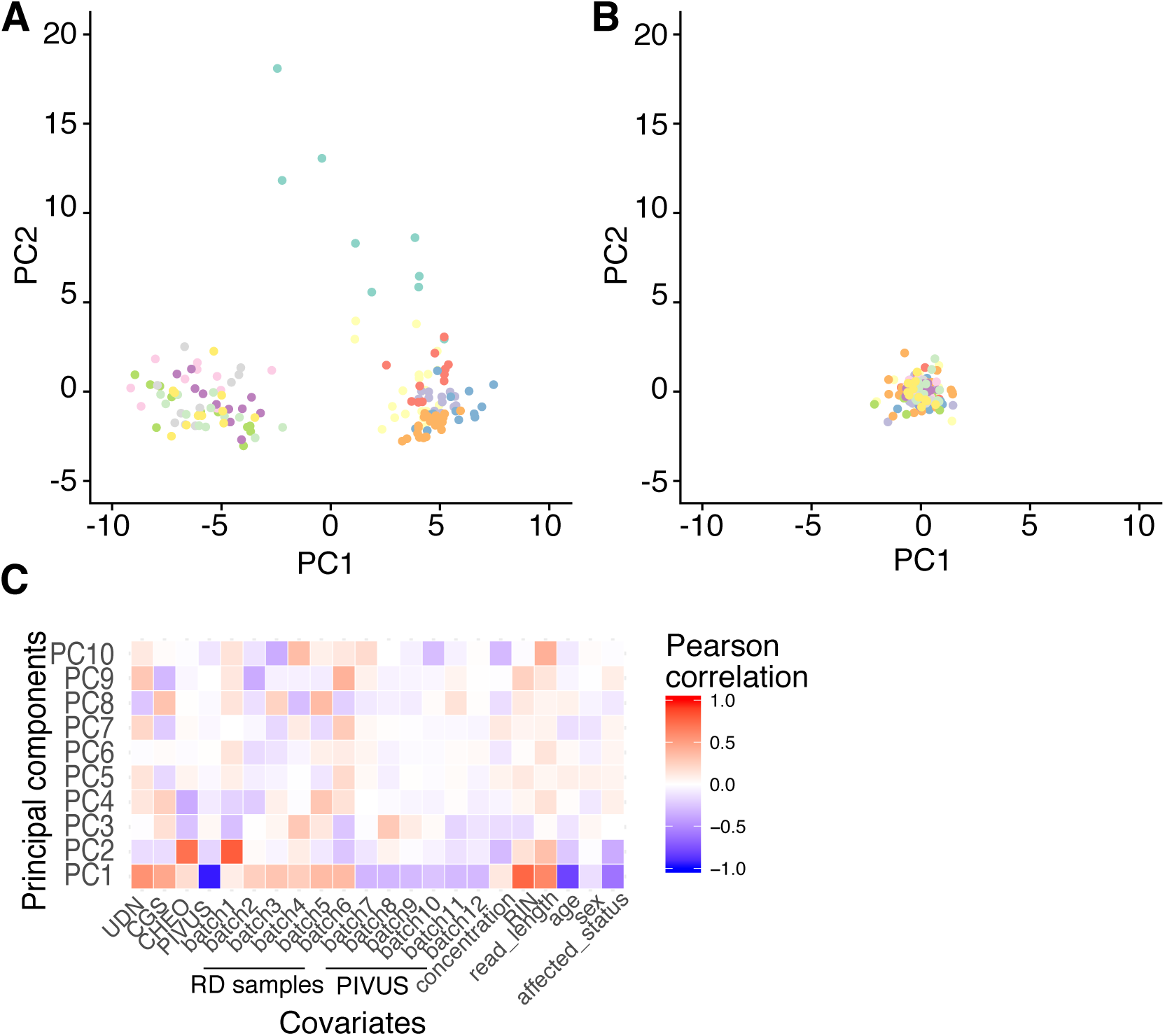
Correction for batch effects - Splicing data. A: Plot of first two principal components (PCs) run on uncorrected splicing ratio data. Samples are colored by batch. We observed that PC1 is separating PIVUS controls samples (left) from rare disease samples (right). B: Plot of first two PCs on splicing ratios after regressing out PCs that explain up to 95% of the variance in the data. Batches are no longer separated on the first PCs. C: Correlation between known covariates 10 first PCs. We do observe that PC1 is highly correlated with the batch, whereas PCs separates samples from one institution (batch 1, CHEO) from others. We also observed that PC1 is highly correlated with RIN, highlighting differences in quality across samples. Overall, the first PCs are correcting for batch effects and quality across our samples.

**Fig. S10:**
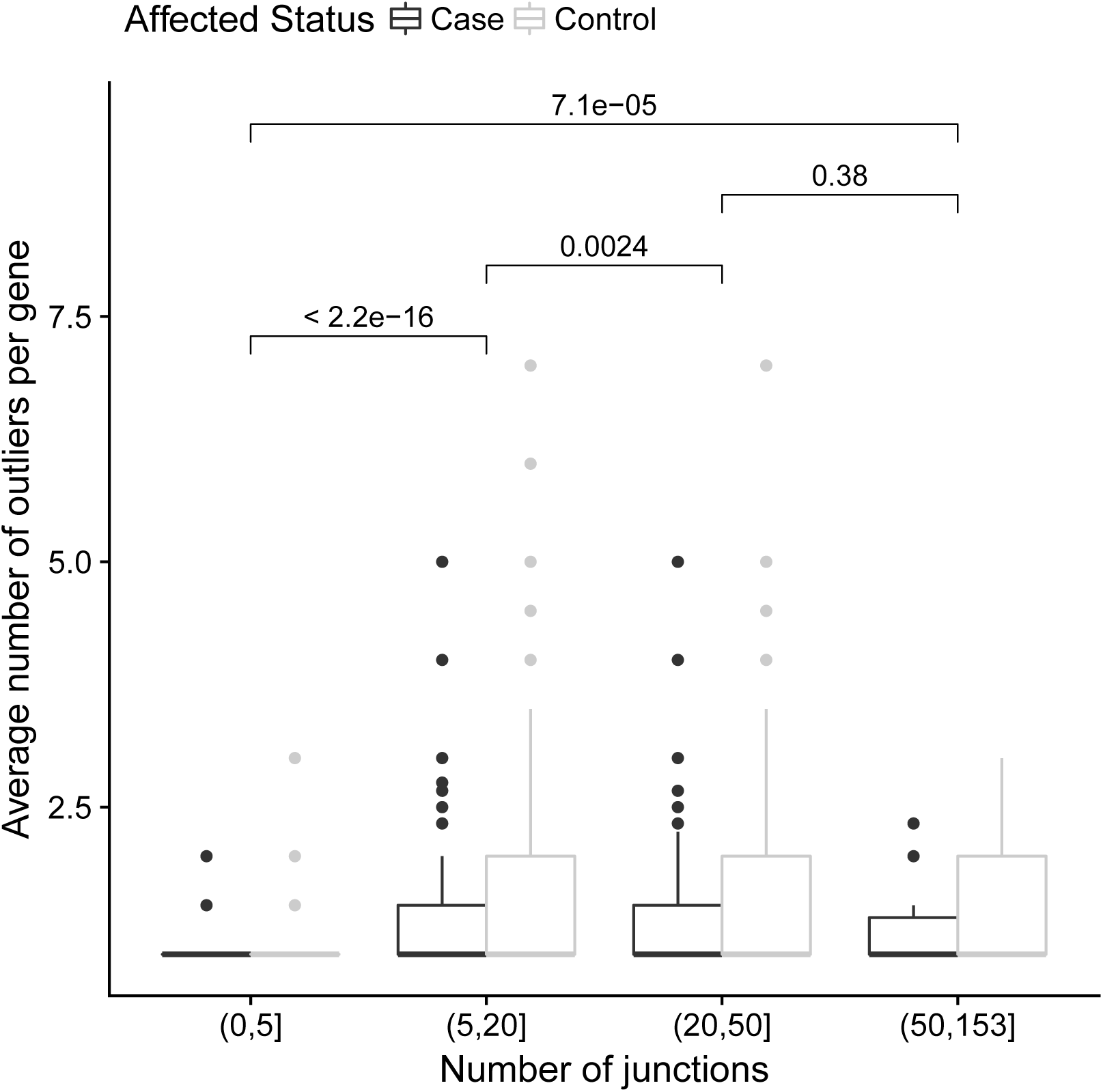
Number of junctions effect on the average number of outliers. We see a significant increase in the average number of outliers in genes when those genes have more splicing junctions (p-values from Wilcoxon test).

**Fig. S11:**
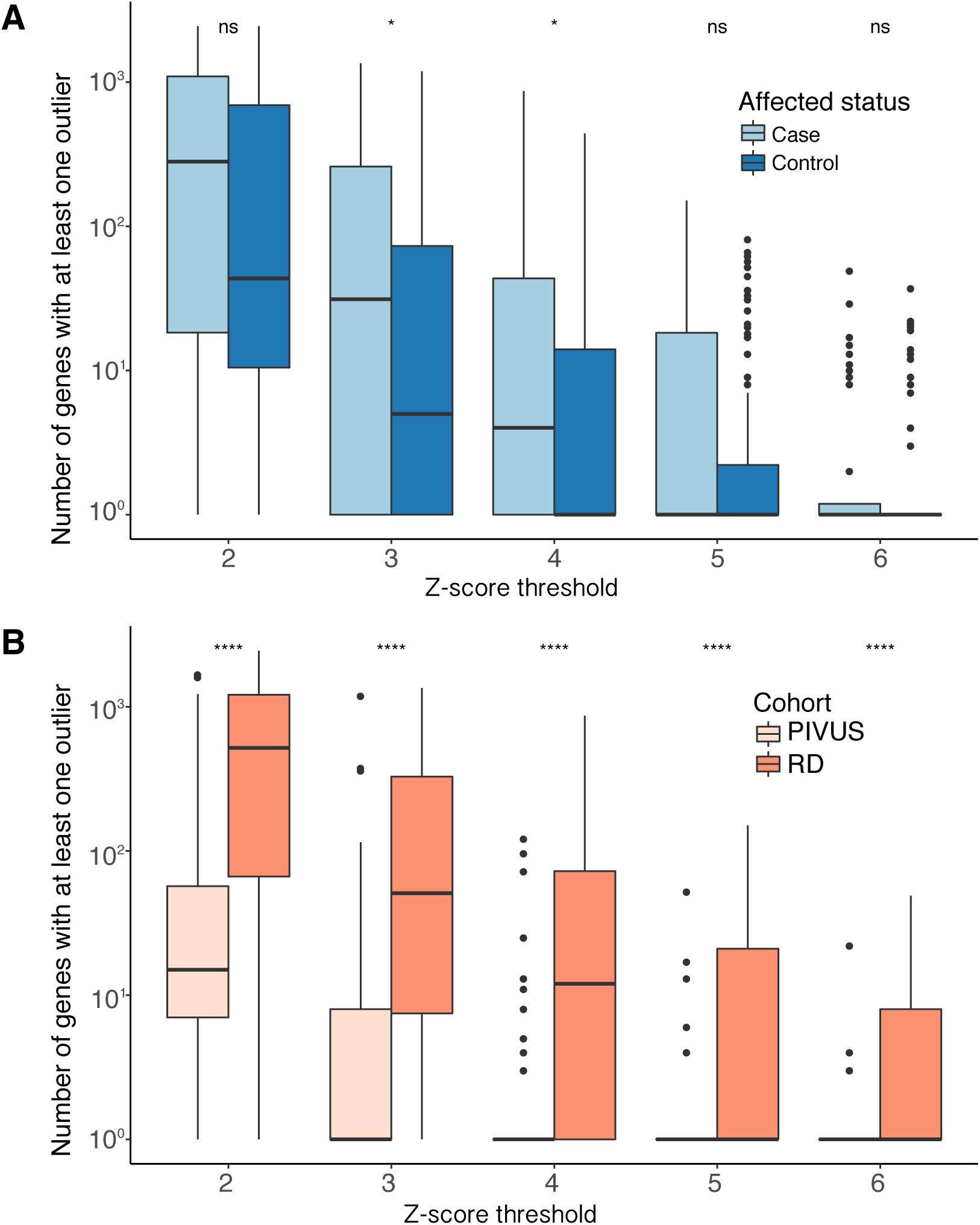
Affected status and cohort effect on splicing Z-score. A: Differences in number of outliers depending on affected status. No significant difference was observed between cases and controls. B: Differences in number of outliers between cohorts. RD: Rare disease. We observed significantly more outliers in the rare disease cohort in comparison to PIVUS (Wilcoxon test).

**Fig. S12:**
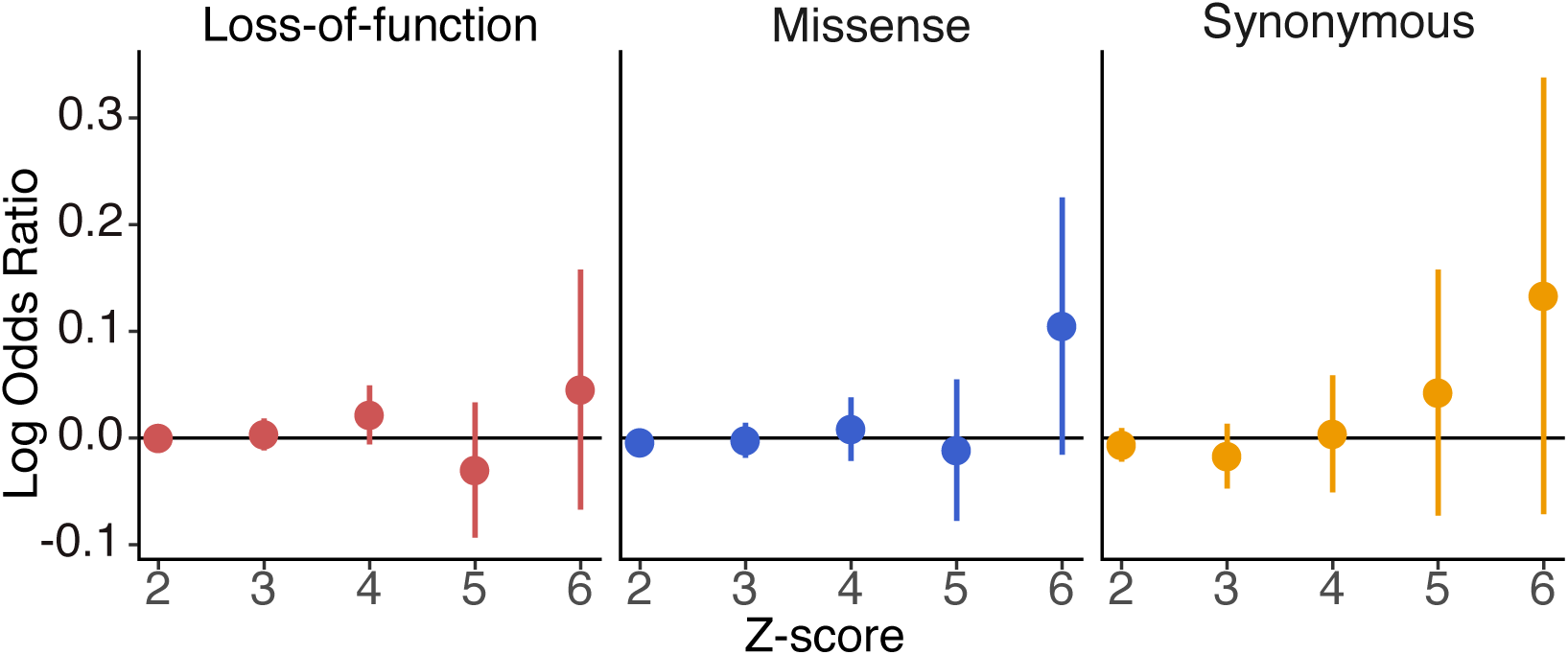
Enrichment patterns for ExAC scores in splicing outliers. We looked at enrichment for cases outlier in genes sensitive to LoF (red), missense (blue) or synonymous variants (yellow) using splicing data. We did not observe a significant enrichment for LoF Z-scores as we do for under-expression outliers. A total of 65 PIVUS were used for controls in this analysis. Due to a lower sample size, those observations can be due to a lack of power or to the intrinsic properties of splicing outliers in opposition to expression outliers.

**Fig. S13:**
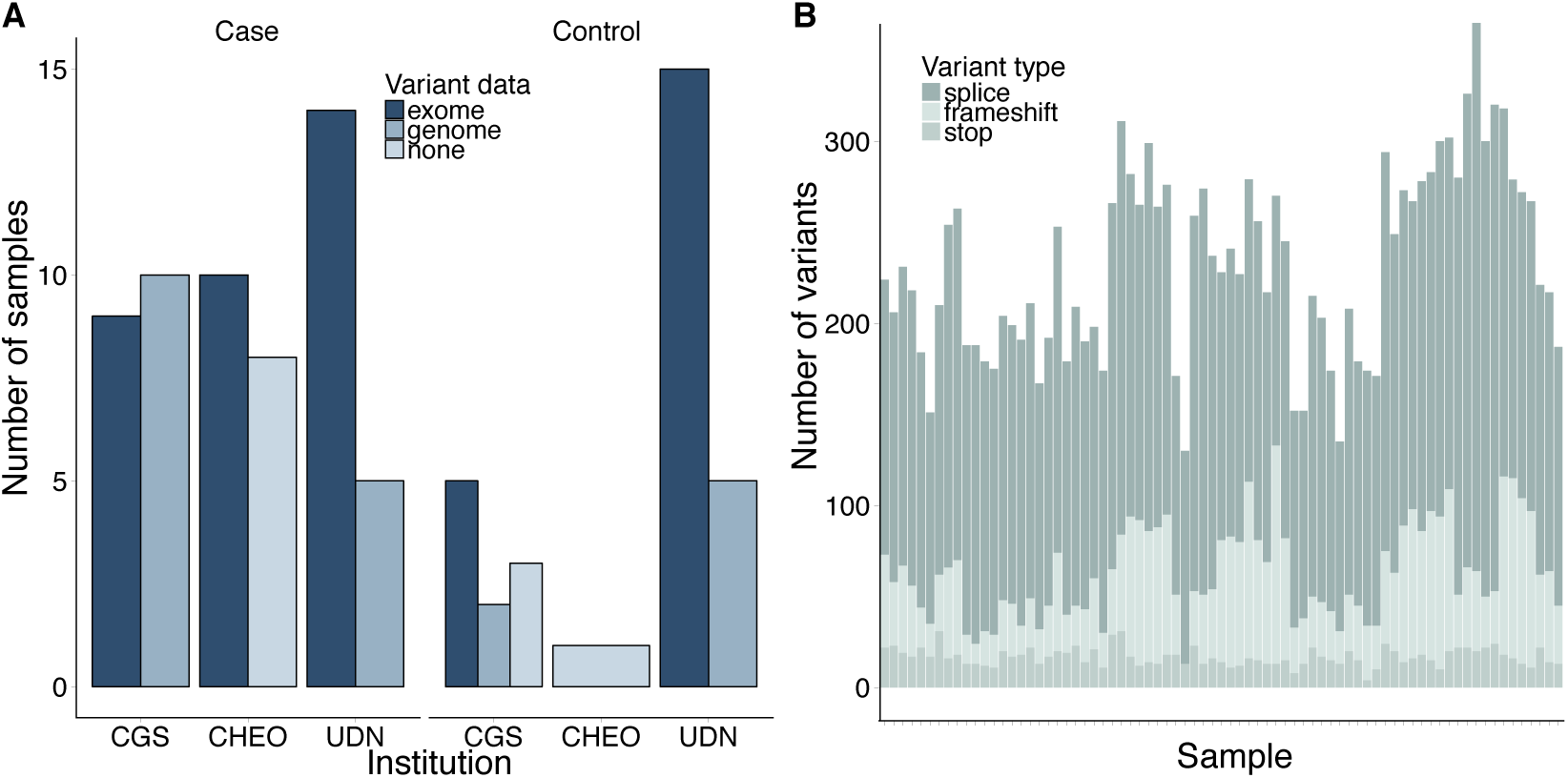
Genetic data across rare disease samples. A: Variant information across rare disease samples and their unaffected family members. We obtained genetic data for 75 samples. B: Number of rare LoF variants across the rare disease samples. Overall, we observed a comparable number of LoF variants across samples.

**Fig. S14:**
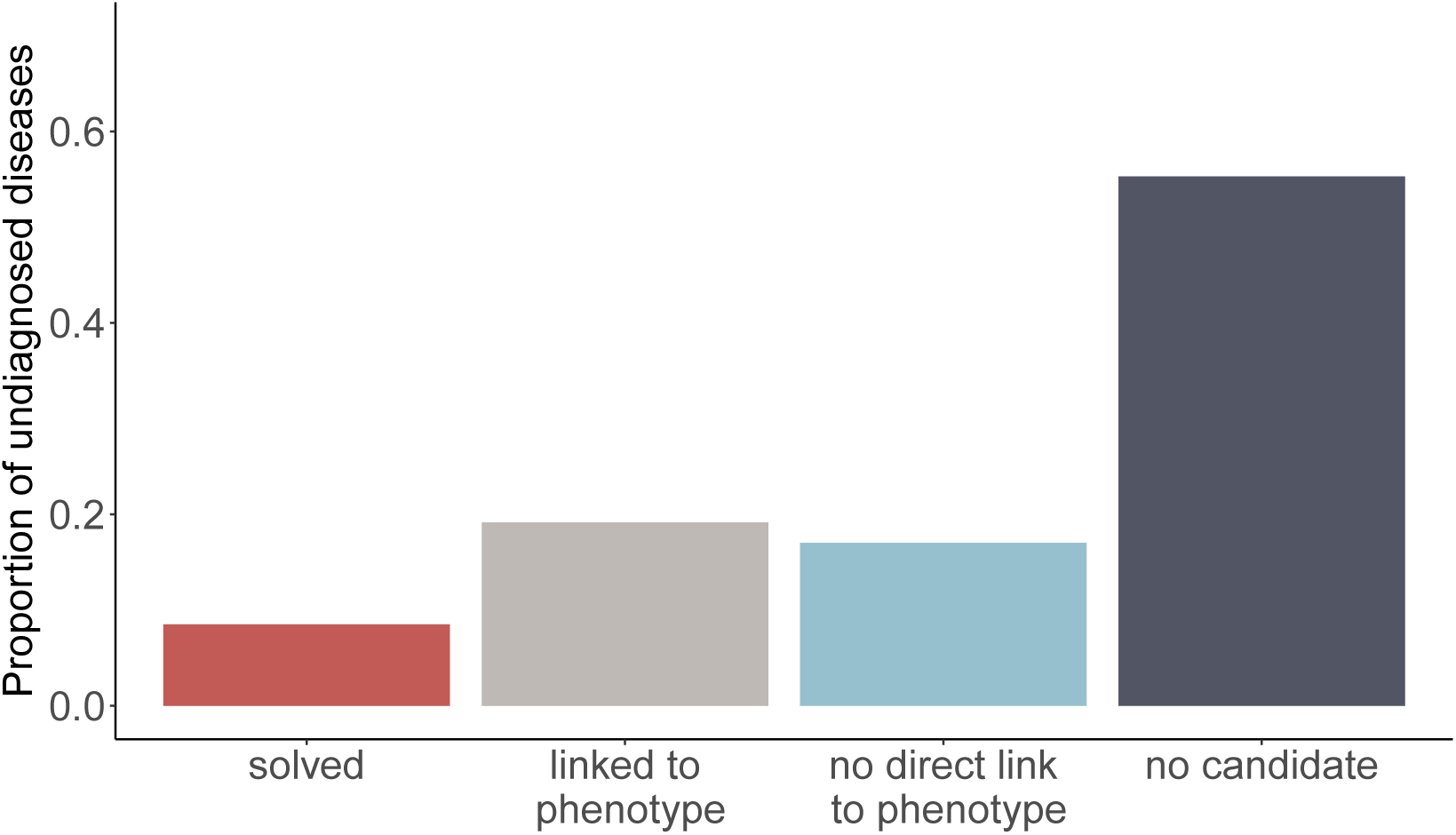
Diagnostic rate after analysis of 47 distinct cases. Solved: causal gene found and further validated. Linked: a gene relevant to the phenotype was found. Not linked: a gene not directly relevant to the phenotype was found. No candidate: no good candidate gene was found

**Fig. S15:**
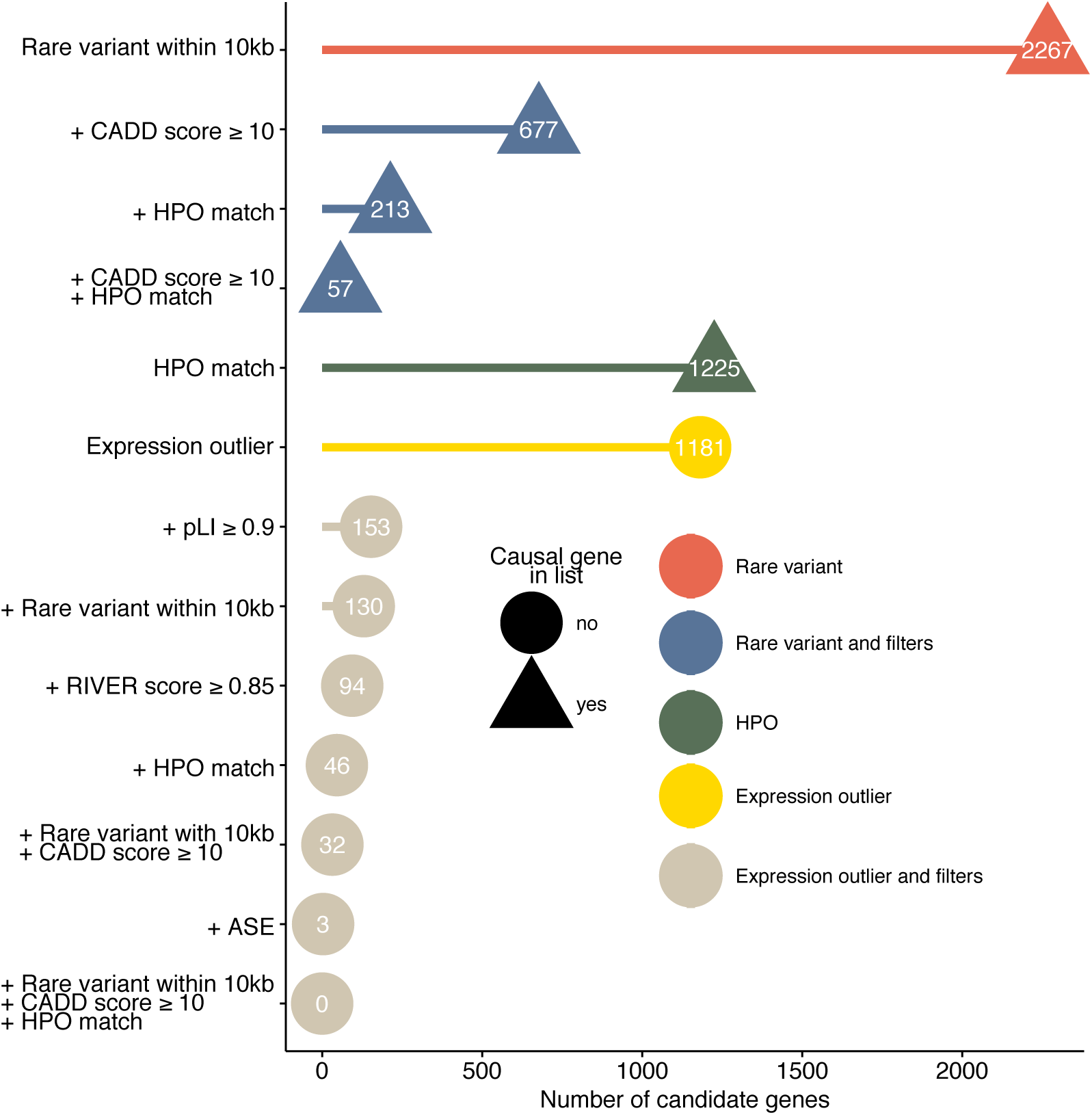
Candidates filters for sister of proband in *RARS2* case. For this sample, *RARS2* Z-score did not pass the |2| threshold (Z-score=-1.55), so the gene was not selected in subsequent filters. However, when we lowered the Z-score threshold to |1.5|, *RARS2* is a candidate among 6 left after filtering the data for deleterious rare variant within 10kb and selecting genes with HPO match.

**Fig. S16:**
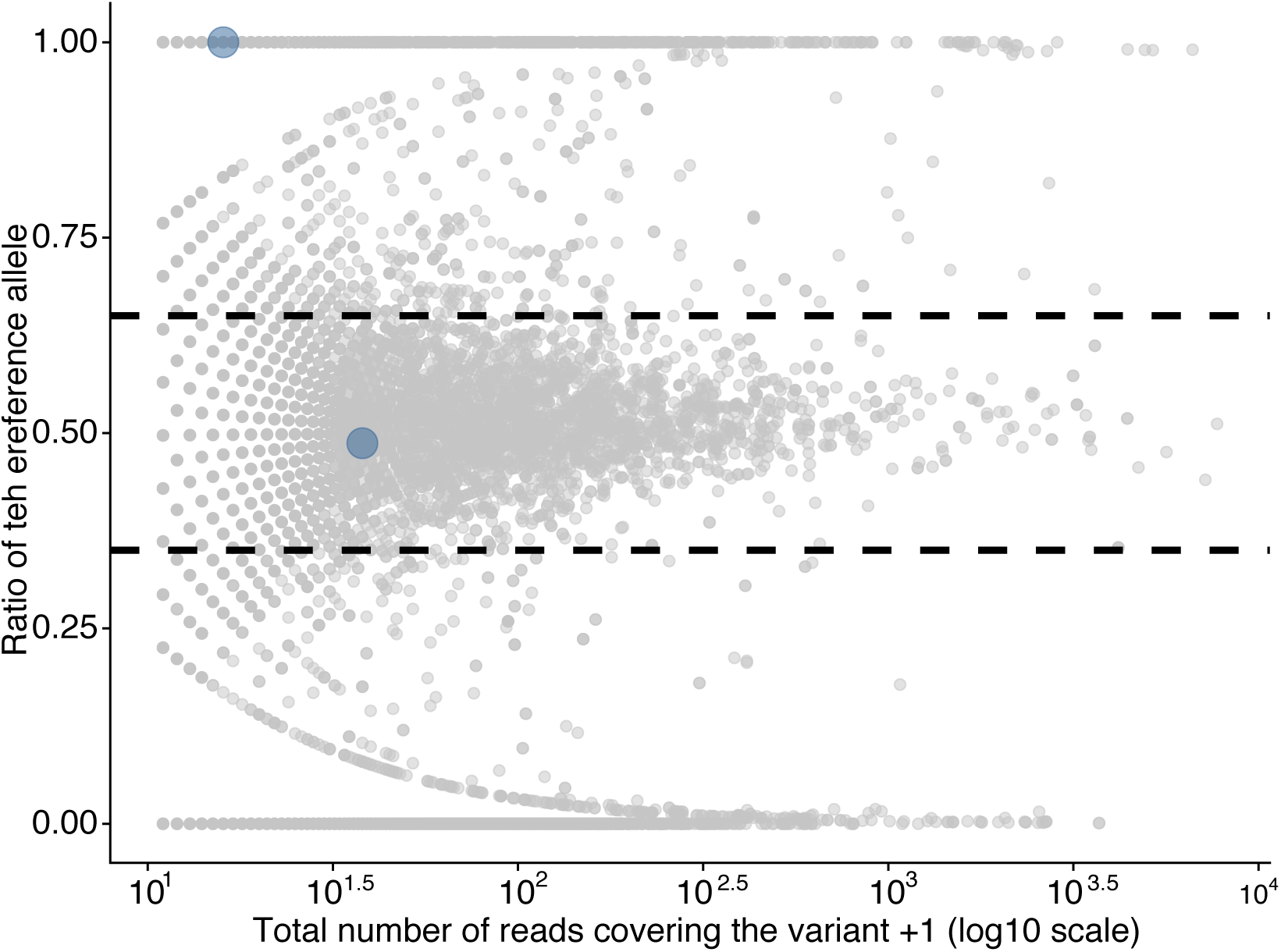
Allelic ratios of variants from second presented case. In blue are variants in *KCTD7* gene. The variant with reference allelic ratio of one corresponds to the causal variant p.V152V. Due to the newly created premature splice junction, only the reference allele is observed.

**Fig. S17:**
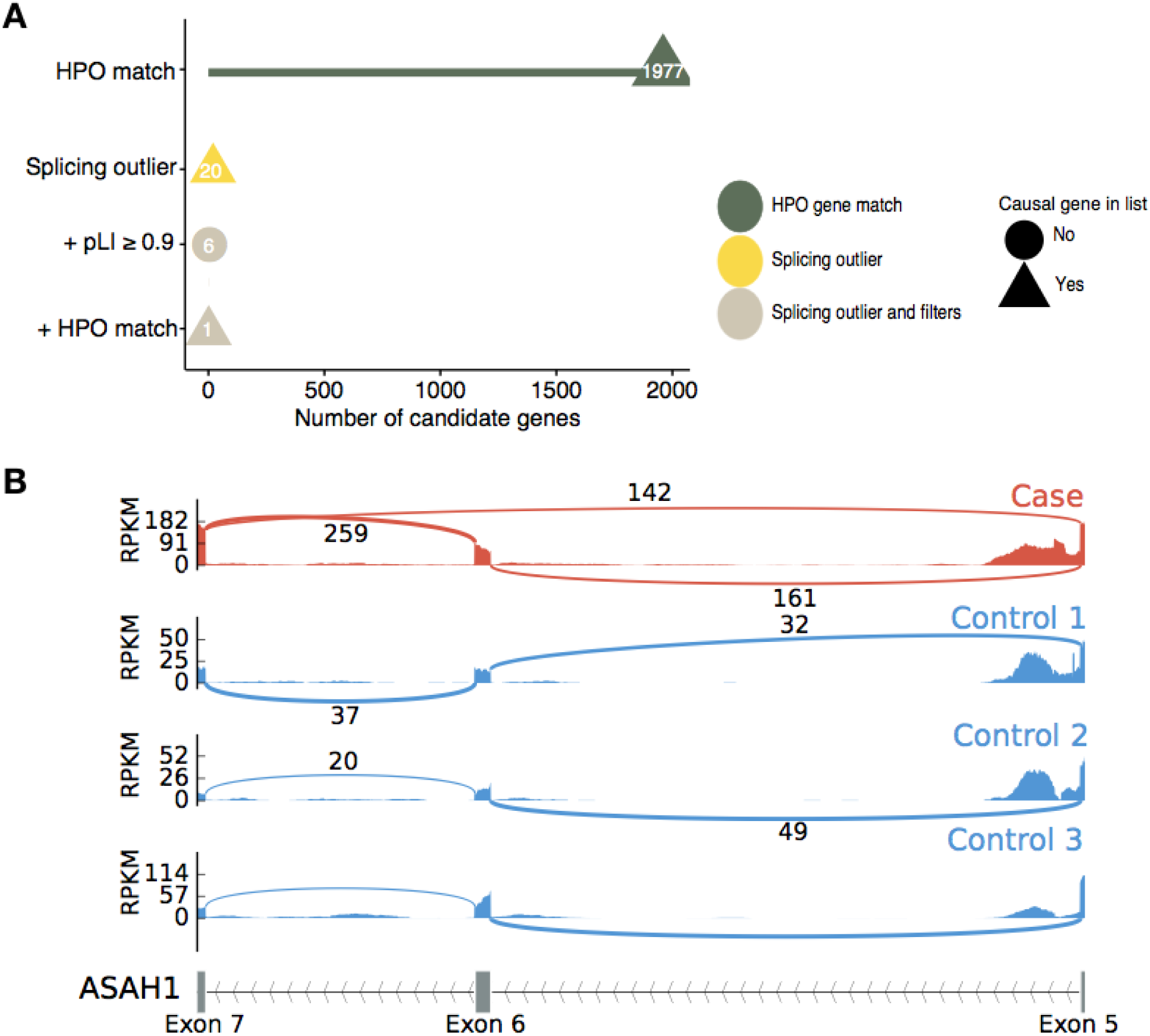
Solved case without genetic data. Example of *ASAH1* gene. A: After filtering our detected splicing outliers for genes related to the phenotype (through HPO Ids), only one candidate was left, *ASAH1*, for which we previously confirmed the association with SMA-PME phenotype in the case. B: Sashimi plot of the case and 2 controls of the *ASAH1* gene. For the case (red track), we observed an alternative transcripts skipping exon 6 (supported by 142 reads). This pattern was never observed in controls.

